# Yeast Ded1 promotes 48S translation pre-initiation complex assembly in an mRNA-specific and eIF4F-dependent manner

**DOI:** 10.1101/353078

**Authors:** Neha Gupta, Jon R. Lorsch, Alan G. Hinnebusch

## Abstract

DEAD-box RNA helicase Ded1 is thought to resolve secondary structures in mRNA 5′-untranslated regions (5′-UTRs) that impede 48S preinitiation complex (PIC) formation at the initiation codon. We reconstituted Ded1 acceleration of 48S PIC assembly on native mRNAs in a pure system, and recapitulated increased Ded1-dependence of mRNAs that are Ded1-hyperdependent in vivo. Stem-loop (SL) structures in 5′-UTRs of native and synthetic mRNAs increased the Ded1 requirement to overcome their intrinsically low rates of 48S PIC recruitment. Ded1 acceleration of 48S assembly was greater in the presence of eIF4F, and domains mediating one or more Ded1 interactions with eIF4G or helicase eIF4A were required for efficient recruitment of all mRNAs; however, the relative importance of particular Ded1 and eIF4G domains were distinct for each mRNA. Our results account for the Ded1 hyper-dependence of mRNAs with structure-prone 5′-UTRs, and implicate an eIF4E·eIF4G·eIF4A·Ded1 complex in accelerating 48S PIC assembly on native mRNAs.

## Introduction

In canonical translation initiation in eukaryotes, a ternary complex (TC), consisting of eukaryotic initiation factor 2 (eIF2), Met-tRNA_i_^Met^, and GTP, along with eIFl, eIFlA, eIF5, eIF4B, and eIF3, binds to the small (40S) ribosomal subunit to form a 43S pre-initiation complex (PIC). The 43S PIC binds to the 5′-end of mRNA and scans the 5′-untranslated region (UTR) to identify the start codon, resulting in the formation of the 48S PIC. eIF4F complex, comprised of eIF4E (a cap binding protein), eIF4G (a scaffolding protein), and eIF4A (DEAD-box RNA helicase), interacts with the mRNA m^7^G cap and aids in recruitment of the 43S PIC to the 5′-end of the mRNA (reviewed in (Dever et al., 2016; Hinnebusch, 2014)).

eIF4A promotes 48S PIC formation in vitro and translation in vivo of virtually all mRNAs regardless of their structural complexity (Pestova and Kolupaeva, 2002; Sen et al., 2015; Yourik et al., 2017). Yeast eIF4A is a weak helicase when unwinding RNA duplexes in vitro (Rajagopal et al., 2012; Rogers et al., 1999). Recent evidence suggests that mammalian eIF4A modulates the structure of the 40S subunit to enhance PIC attachment (Sokabe and Fraser, 2017). Ded1 is a yeast DEAD-box RNA helicase that promotes translation in vivo of reporter mRNAs with longer or structured 5′-UTRs (Berthelot et al., 2004; Chiu et al., 2010; Sen et al., 2015). Like eIF4A, Ded1 is essential for yeast growth and stimulates bulk translation initiation in vivo (Chuang et al., 1997; de la Cruz et al., 1997). However, ribosome profiling of conditional *Ded1* mutants revealed that native mRNAs with 5′-UTRs that are longer and more structured than average yeast 5′-UTRs exhibit a greater than average reduction in translational efficiency (TE) on Ded1 inactivation (Ded1-hyperdependent mRNAs); whereas mRNAs with shorter and less structured 5′-UTRs exhibit increased relative TEs in *ded1* cells (Ded1-hypodependent mRNAs) (Sen et al., 2015).

Yeast Ded1 can unwind model RNA duplexes and act as an RNA chaperone or RNA-protein complex remodeler in vitro (Iost et al., 1999; Bowers et al., 2006; Yang and Jankowsky, 2006). Ded1 can physically interact individually with eIF4E, eIF4G, and eIF4A, as well as with eIF4A and eIF4G simultaneously in an RNA-independent manner (Hilliker et al., 2011; Senissar et al., 2014; Gao et al., 2016). Ded1 interacts with eIF4A through its N-terminal domain (Ded1-NTD) and this interaction is required for stimulation of Ded1’s RNA-duplex unwinding activity by eIF4A (Gao et al., 2016). Yeast eIF4G contains three RNA binding domains, N-terminal RNA1, central RNA2, and C-terminal RNA3 (Berset et al., 2003); and while none of the three is essential, simultaneous deletion of RNA2 and RNA3 is lethal (Park et al., 2011). In vitro, eIF4G variants lacking any of the three RNA binding domains exhibit similar affinities for eIF4A, support similar rates of ATP-hydrolysis by the eIF4F complex (albeit with higher K_m_ values for ATP), but lack the preference of WT eIF4F for 5′-overhang substrates during unwinding (Rajagopal et al., 2012). The Ded1 C-terminal domain (Ded1-CTD) interacts with the eIF4G-RNA3 domain, and Ded1-eIF4G interaction decreases the rate of RNA unwinding while increasing Ded1 affinity for RNA in vitro (Hilliker et al., 2011; Putnam et al., 2015). Ded1 also interacts with the RNA2 domain of eIF4G and with eIF4E (Senissar et al., 2014), but the physiological relevance of these interactions are unknown.

We previously demonstrated functions of eIF4F, eIF4B, and eIF3 in stimulating the rate and extent of mRNA recruitment by 43S PICs in a fully purified yeast system for native *RPL41A* mRNA, containing a short and relatively unstructured 5′-UTR (Mitchell et al., 2010). Although Ded1 is essential in vivo, it was dispensable for recruitment of this mRNA in vitro. Considering that *RPL41A* was judged to be Ded1-hypodependent in vivo by ribosome profiling of *ded1* mutants (Sen et al., 2015), we asked whether recruitment of Ded1-hyperdependent mRNAs would require Ded1 in the reconstituted system. We investigated whether the presence of defined stem-loop (SL) structures in native or synthetic mRNAs would confer greater Ded1-dependence for rapid recruitment in vitro. Finally, we examined the role of the RNA2 and RNA3 domains of eIF4G and the NTD and CTD of Ded1 that mediate Ded1 interactions with the eIF4F complex in promoting Ded1’s ability to accelerate mRNA recruitment by 43S PICs. Our findings demonstrate that Ded1 accelerates recruitment of native and synthetic mRNAs, overcoming the inhibitory effects of structured leader sequences and conferring relatively greater stimulation for mRNAs hyperdependent on Ded1 in vivo, all in a manner consistent with stimulation by formation of a Ded1-eIF4F complex.

## Results

### Ded1 enhances the rate of recruitment of all natural mRNAs tested

We set out to reconstitute the function of Ded1 in 48S PIC assembly in a yeast translation initiation system comprised of purified components (Mitchell et al., 2010; Walker et al., 2013; Yourik et al., 2017). Pre-assembled 43S PICs, containing 40S subunits and factors eIF1, eIF1A, eIF5, eIF2-GDPNP-Met-tRNA_i_, eIF4·G4E, eIF4A, eIF4B, and eIF3, were pre-incubated with or without Ded1, and reactions were initiated by addition of ATP and ^32^P-labeled m^7^Gppp-capped mRNA (synthesized in vitro). Formation of 48S complexes was monitored over time using a native gel electrophoretic mobility shift assay (EMSA) to resolve free and 48S-bound mRNAs. An ~20-fold excess of unlabeled-capped mRNA was added to reaction aliquots at each time point to quench further recruitment of ^32^P-labeled mRNA (“pulse-quench”). By varying the concentration of Ded1, this assay yields the apparent rate constants (k_app_) for 48S PIC formation at each Ded1 concentration, the maximal rate at saturating Ded1 (k_max_), the Ded1 concentration required for the half-maximal rate of 48S formation (K_1/2_), and the reaction endpoints (percentage of mRNA recruited) at each Ded1 concentration. The addition of non-radiolabeled mRNA in the quench did not dissociate the pre-formed ^32^P-labeled 48S complexes (Figs. S1A, S2E). Our purified Ded1 hydrolyzed ATP in an RNA-dependent manner with k_cat_ and 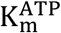 values consistent with previous measurements (Figs. S1B-D) (Iost et al., 1999; Senissar et al., 2014). Moreover, the Ded1 bound a fluorescently labeled single-stranded mRNA in the presence or absence of ADP or ADPNP with K_D_ values consistent with published values (Fig. S1E) (Banroques et al., 2008; Iost et al., 1999).

We began by analyzing the effect of Ded1 on recruitment of *RPL41A* mRNA, a short transcript of 310 nucleotides (nt), with 5′-UTR of only 24 nt (Figs. 1A, S1F), and a low degree of predicted secondary structure (Mitchell et al., 2010). *RPL41A* behaved like a Ded1-hypodependent mRNA in ribosome profiling experiments, exhibiting increased relative TE in *ded1* mutant versus WT cells (Sen et al., 2015). Consistent with this, recruitment of *RPL41A* mRNA in vitro was achieved previously without Ded1 (Mitchell et al., 2010). Nevertheless, with our more sensitive pulse-quench approach, we found that addition of saturating Ded1 increased the k_max_ of *RPL41A* by ~2.8-fold from 0.95 ± 0.1 min^−1^ to 2.7 ± 0.3 min^−1^ (Figs. 1B, cf. blue vs. orange, & S1G-red), with a 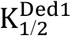 of 58 ± 8 nM (Fig. 1C). Thus, despite its low Ded1-dependence relative to most other mRNAs in vivo, *RPL41A* recruitment is appreciably stimulated by Ded1 in vitro.

We next examined Ded1 stimulation of another Ded1-hypodependent mRNA, *HOR7*, and several Ded1-hyperdependent mRNAs, *SFT2, PMA1, OST3*, and *FET3*, which exhibited reduced TEs in *ded1* versus WT cells (Sen et al., 2015). *SFT2* and *PMA1* mRNAs are noteworthy in containing SL structures in their 5′-UTRs detected in vivo (Rouskin et al., 2014). Because these mRNAs exceed the maximum length that can be resolved by EMSA (~400 nt), we generated reporter mRNAs with the 5′-UTR and first 60 nt of coding sequences (CDS) of each mRNA (Figs. 1A, S1F). (For brevity, we refer to these reporter mRNAs simply by their gene names.) Without Ded1, the rate of *HOR7* recruitment was 1.3 ± 0.l6 min^−1^, and saturating Ded1 stimulated recruitment by ~2-fold (k_max_ = 2.8 ± 0.4 min^−1^) with a 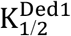 of 115.5 ± 17 nM, results that were similar to those for Ded1-hypodependent *RPL41A* (Figs. 1B, 1C, S1G-blue curve). Strikingly, without Ded1, the four Ded1-hyperdependent mRNAs were recruited at rates 2.5-to 15-fold lower compared to the two Ded1-hypodependent mRNAs (Fig. 1B, blue), and these rates increased by an order of magnitude on addition of Ded1 (Figs. 1B, orange & bottom panel; S1G). The Ded1-hyperdependent mRNAs also required higher Ded1 concentrations to achieve these maximal rates (Fig. 1C). The acceleration of mRNA recruitment by Ded1 required its ATPase activity, as ATPase-deficient Ded1 variant Ded1^E307A^ (Figs. S1C-D) (Iost et al., 1999) did not accelerate recruitment of any mRNAs (Fig. S1H).

**Fig. 1:**
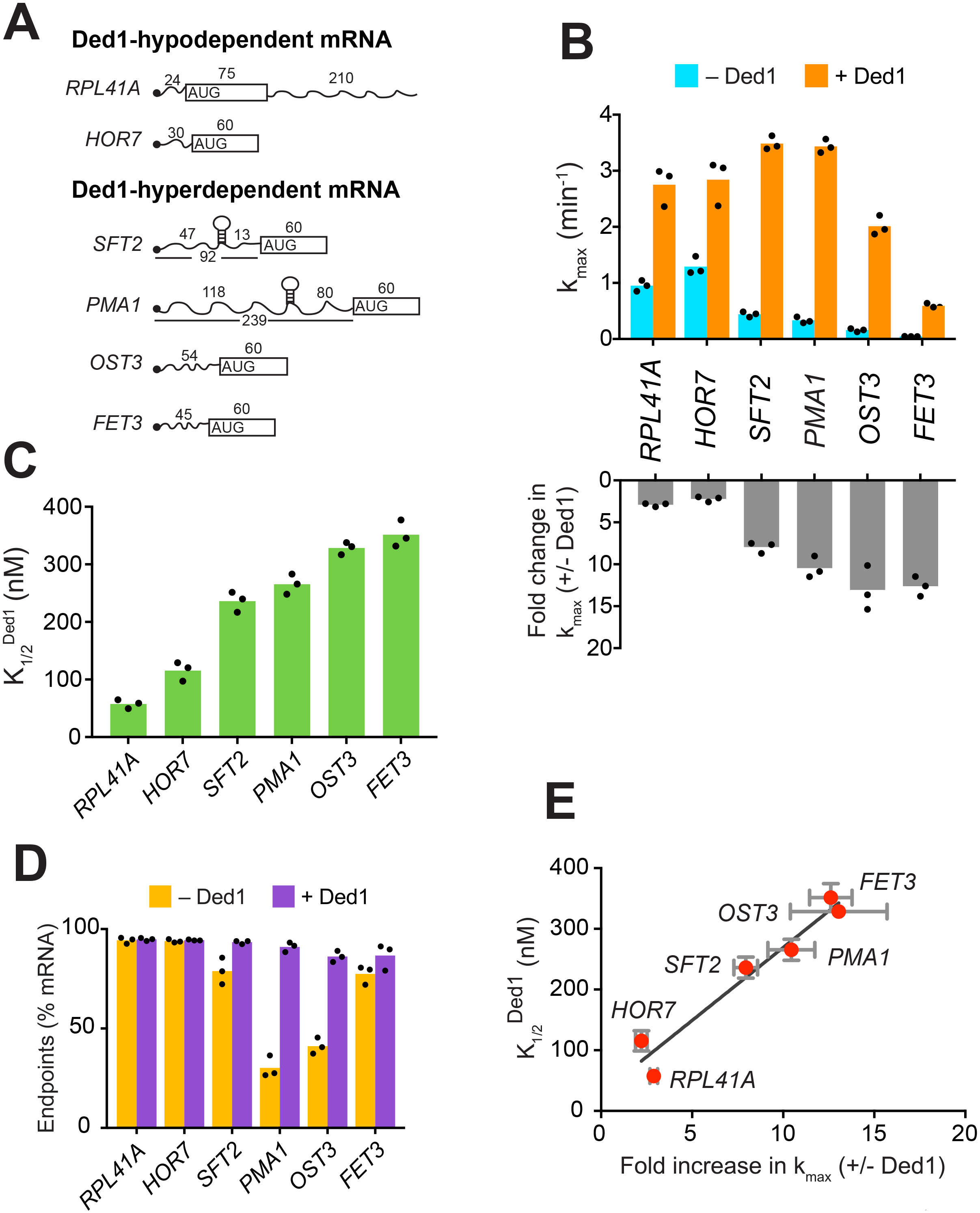
Ded1 confers relatively greater acceleration of 48S PIC assembly in vitro for native mRNAs hyperdependent on Ded1 in vivo. **(A)** Schematics of native *RPL41A* mRNA, and mRNA reporters for other native yeast mRNAs comprised of the 5′-UTR and first 60 nt of the ORF. ORFs are depicted as boxes with an AUG start codon; wavy lines depict 5′- and 3′-UTRs of the indicated nucleotide lengths; black balls depict m^7^Gppp caps; SLs in the 5′-UTRs of *SFT2* and *PMA1* identified in vivo are shown as hairpins, whose folding free energies are given in Fig. S2A. (B) *Upper:* Maximum rates of recruitment (k_max_) in the absence (blue) and presence (orange) of saturating Ded1 for mRNAs in (A). *Lower:* Fold-changes in k_max_ with or without Ded1 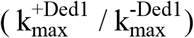 calculated from data in upper plot. **(C)** Concentrations of Ded1 resulting in half-maximal rates 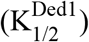. (D) Endpoints of mRNA recruitment as percentages of total mRNA (15 nM) bound to 48S PICs (30 nM) in absence (gold) or presence (purple) of saturating Ded1. (E) Plot of 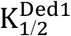 from (C) versus fold-change in 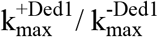 from (B, *lower*) for each mRNA. Red points and error bars indicate mean values and one standard deviation, respectively, for each parameter. Line generated by linear regression analysis (Y = 24.02*X + 28.98, R^2^ = 0.944, P-value = 0.001). (B-D) Bars indicate mean values calculated from the 3 independent experiments represented by the black data points. See Supplementary Figures 1 and 2 and Figure1_SourceData1

*PMA1* and *OST3* also exhibited low endpoints of recruitment without Ded1, 30 ± 5% and 41 ± 4%, respectively, which increased to 91 ± 2% and 86 ± 3%, respectively, on Ded1 addition (Fig. 1D). It was suggested that Ded1 acts as an RNA chaperone to aid transitions between different RNA conformations (Yang et al., 2007). In fact, two or more conformers of *OST3* (and other mRNAs) were observed in native gel electrophoresis that likely represent differently folded, stable mRNA conformers, and Ded1 resolved each of them into one major species (Fig. S1I). Perhaps only one of these conformers of *PMA1* and *OST3* is competent for 48S PIC assembly, and Ded1 facilitates isomerization among them.

Interestingly, a linear relationship was observed between the fold-acceleration by Ded1 and the 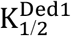, such that Ded1-hypodependent and Ded1-hyperdependent mRNAs cluster separately from each other along the line (Fig. 1E). One explanation could be that Ded1-hyperdependent mRNAs have a higher rate-limiting activation energy barrier for 48S PIC assembly in the absence of Ded1 compared to the Ded1-hypodependent mRNAs, consistent with the latter’s relatively higher rates of recruitment without Ded1 (Fig. 1B, blue bars, *RPL41A* and *HOR7* versus *SFT2, PMA1, OST3*, and *FET3*). Accordingly, the hyperdependent mRNAs require relatively higher Ded1 concentrations to lower this barrier to the point at which a Ded1-independent step becomes rate limiting *(SFT2* and *PMA1)* or where Ded1 cannot lower the Ded1-dependent barrier further *(OST3* and *FET3)* (Fig. S1J).

Taken together, the data presented above demonstrate that mRNAs with long and structured 5′-UTRs, found to be hyperdependent on Ded1 for translation in vivo, are inherently less capable of PIC recruitment and more dependent on Ded1 for rapid recruitment in vitro than are mRNAs hypodependent on Ded1 in vivo.

### Secondary structures in the 5′-UTR increase Ded1-dependence in 48S PIC assembly

Given that mRNAs with heightened Ded1-dependence in vivo have a greater than average potential to adopt secondary structures involving the 5′-UTR (Sen et al., 2015), we investigated if the stable SL structures previously detected in the 5′-UTRs of *SFT2* and *PMA1* were responsible for their elevated Ded1 dependence (Rouskin et al., 2014). To this end, we introduced mutations to eliminate the SL in each mRNA, or (for *SFT2*) to strengthen the SL (Figs. 2A, S2A-B). Without Ded1, the SL-disrupted version of *PMA1*, *PMA1-M*, showed ~2-fold higher endpoints of recruitment than WT *PMA1* (*PMA1-M* = 62 ± 2.5%, *PMA1* = 30 ± 5%, Fig. 2B, gold). *PMA1-M* was also recruited at rates ~4-fold higher than *PMA1* (k_max_ = 1.2 ± 0.1 min^−1^ *(PMA1-M*) vs. 0.33 ± 0.06 min^−1^ (*PMA1*); Fig. 2C, blue), consistent with the idea that the SL inhibits mRNA recruitment. Ded1 increased the rate of *PMA1-M* recruitment to yield a k_max_ similar to that for *PMA1* (Fig. 2C, orange), but at a much lower Ded1 concentration for *PMA1-M* 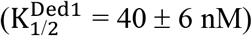 versus *PMA1* (~280 nM) (Fig. 2D; S2C, black vs. purple). Similar results were observed for the SL-disrupted *SFT2* variant, *SFT2-M*, as follows. Compared to WT *SFT2* mRNA, *SFT2-M* showed an ~2-fold faster recruitment in the absence of Ded1 (0.88 ± 0.08 min^−1^ vs. 0.44 ± 0.05 min^−1^ Fig. 2C, blue) and an ~2-fold lower Ded1 concentration required to accelerate recruitment to half the maximum rate: 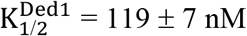 (*SFT2-M*) vs. 236 ± 18 nM (*SFT2*) (Figs. 2C, orange; 2D; S2C, green vs. orange curves). Importantly, both *PMA1-M* and *SFT2-M* mRNAs clustered with the Ded1-hypodependent mRNAs instead of the Ded1-hyperdependent mRNAs in the plot of fold-acceleration versus 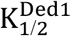 (Fig. 2E, blue). In contrast, the *SFT2-M2* variant harboring a SL of enhanced stability (ΔG° = -19.1 kcal/mol vs. −9.4 kcal/mol for WT *SFT2;* Figs. 2A, S2A, and S2B) was not recruited without Ded1, and exhibited only low-level recruitment at the highest achievable Ded1 concentration (Fig. S2D). These results strongly suggest that Ded1 accelerates the recruitment of WT *SFT2* and *PMA1* mRNAs, and increases the end-point of *PMA1* recruitment, in part by melting the SL structures in their 5′-UTRs. As the *PMA1* SL is ~120 nt from the 5′-cap, it is likely that Ded1 resolves the SL to accelerate scanning of the PIC through the 5′-UTR.

**Fig. 2:**
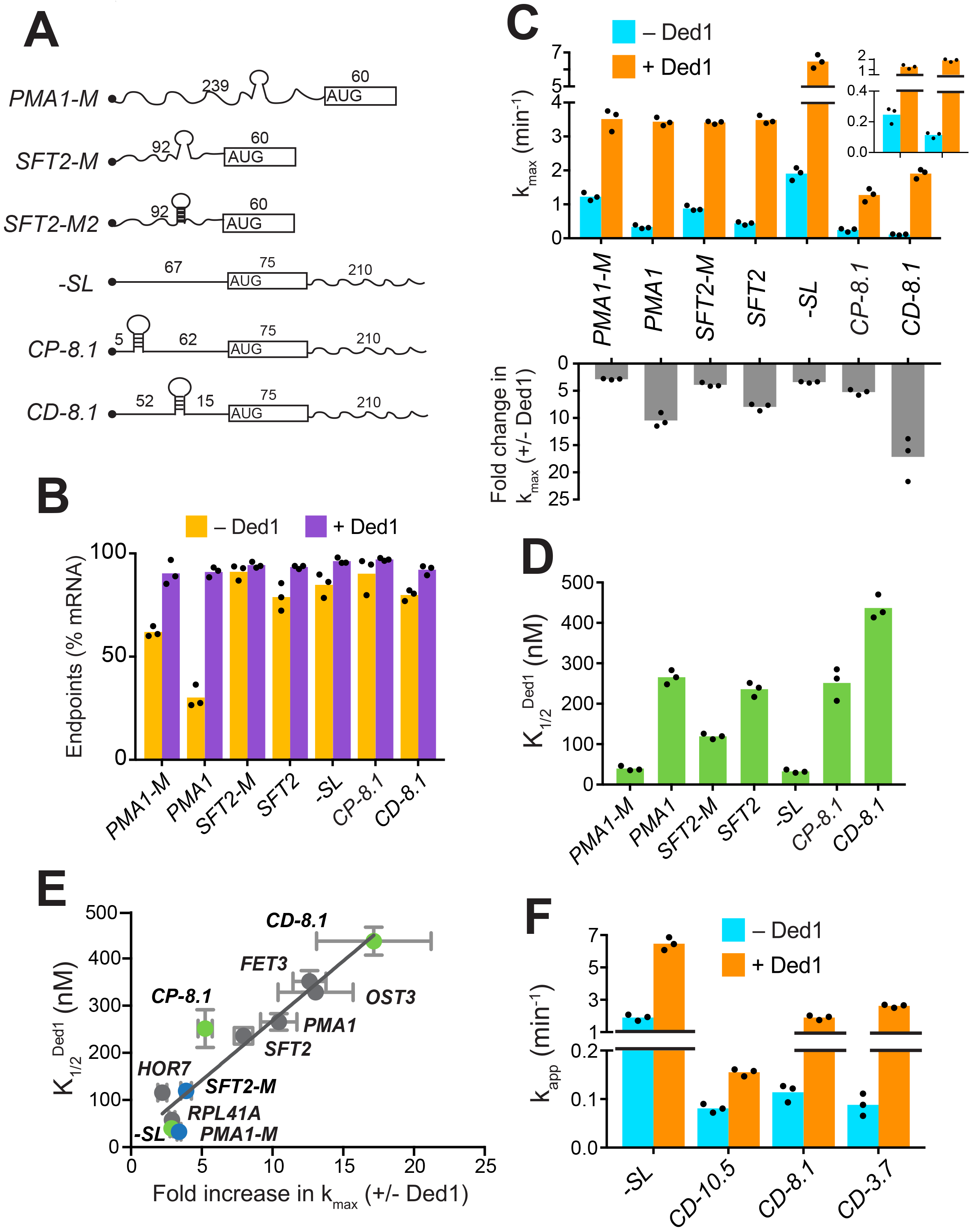
Secondary structures in 5′-UTRs confer Ded1-hyperdependence in accelerating 48S PIC assembly in vitro. **(A)** Schematics of derivatives of natural mRNAs from Fig. 1A mutated to decrease (*SFT2-M* and *PMA1-M*) or increase (*SFT2-M2*) the stability of 5′-UTR SLs; and synthetic mRNAs depicted as in Fig. 1A. The sequences of SLs and mutations are in Fig. S2A. **(B)** Endpoints of 48S PIC assembly reactions, determined as in Fig. 1D. **(C)** k_max_ values (*upper*) and fold-change in k_max_ (*lower*) in the presence versus absence of Ded1, determined as in Fig. 1B. The inset in the upper panel scales the y-axis to display the low k_max_ values for *CD-8.1* and *CP-8.1* mRNAs without Ded1. Note that Y-axis is discontinuous. **(D)** 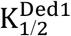 values determined as in Fig. 1C. **(E)** Plot of 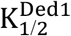 from (D) versus fold-change in 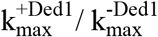 from (C, *lower*) for the indicated mRNAs. Gray, blue, and green points indicate natural, mutated, and synthetic mRNAs, respectively, with error bars indicating 1 SD; line produced by linear regression analysis (Y = 25.29*X + 15.28, R^2^ = 0.89, P-value < 0.001). **(F)** Apparent rate of mRNA recruitment of synthetic mRNAs with different strengths SLs in the cap-distal region in the absence (blue) and presence of Ded1 (orange). AG°: *CD-10.5* = −10.5 kcal/mol, *CD-8.1* = −8.1 kcal/mol, and *CD-3.7* = −3.7 kcal/mol. Note that Y-axis is discontinuous. (B-D, F) Bars indicate mean values calculated from the 3 independent experiments represented by the data points. (B-D) WT *PMA1* amd WT *SFT2* data is added from Fig. 1 for comparison. See Supplementary Figure 2 and Figure2_SourceData1

### Evidence that Ded1 stimulates the PIC attachment and scanning steps of initiation

Although Ded1-hypodependent mRNAs *RPL41A* and *HOR7* lack any strong, defined SLs, Ded1 still stimulated their recruitment (Fig. 1B). Similarly, even after removal of SLs, the *PMA1-M* and *SFT2-M* mutant mRNAs were still stimulated by Ded1 (Fig. 2C). In addition to defined, stable SLs in 5′-UTRs, natural mRNAs likely form large ensembles of weaker structures involving interactions of nucleotides within the 5′-UTR or between 5′-UTR nucleotides and nucleotides in the CDS or 3′-UTR (Kertesz et al., 2010; Yourik et al., 2017), which might also contribute to Ded1-dependence during PIC attachment or scanning. To test this hypothesis, we examined recruitment of chimeric mRNAs with synthetic 5′-UTRs attached to the CDS and 3′-UTR of native *RPL41A* (Yourik et al., 2017). A synthetic mRNA dubbed “-SL” (for “minus stem-loop”) contained a 67 nt 5′-UTR comprised of CAA repeats that is devoid of stable secondary structure (Figs. 2A, S2B). Without Ded1, *-SL* was recruited rapidly at a rate of 1.9 ± 0. 2 min^−1^ (Fig. 2C, blue), ~2-fold faster than *RPL41A* (Fig. 1B, blue) or the *SFT2-M* and *PMA1-M* variants lacking SLs (Fig. 2C, blue), consistent with the idea that weaker interactions formed by 5′-UTRs of native mRNAs lacking SLs can inhibit recruitment. Ded1 increased the k_max_ of *-SL* recruitment by ~ 3-fold (Figs. 2C, orange; S2F, red), comparable to the increases observed for *SFT2-M, PMA1-M* (Fig. 2C, *lower)*, and *RPL41A* (Fig. 1B, *lower*). This stimulation of *-SL* recruitment was achieved at a relatively low Ded1 concentration comparable to, or lower than, those observed for *SFT2-M, PMA1-M*(Fig. 2D), and *RPL41A* (Fig. 1C). Hence, *-SL* mRNA clustered with the Ded1-hypodependent mRNAs shown in the plot of 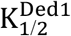 versus fold-increase in k_max_ (Fig. 2E, green, *-SL*). The finding that Ded1 significantly stimulates 48S PIC assembly on *-SL*, containing a 5′-UTR incapable of forming stable structures on its own, suggests that Ded1 has a second role in mRNA recruitment, in addition to unwinding stable structures in 5′-UTRs.

To analyze whether Ded1 can stimulate the PIC attachment step of 48S assembly, we examined the synthetic *CP-8.1* mRNA with a stable SL (predicted ΔG° = −8.1 kcal/mol) inserted in a cap-proximal location 5 nt from the 5′-end of *-SL* mRNA. We also analyzed a third synthetic mRNA, *CD-8.1*, containing the same SL inserted 45 nt downstream from the cap of *-SL*, reasoning that this cap-distal SL might impede scanning without interfering with PIC attachment at the cap (Figs. 2A, S2A, S2B). As expected, both synthetic mRNAs with SLs had order-of-magnitude lower rates of recruitment compared to *-SL* in the absence of Ded1 (Fig. 2C, blue, inset), indicating that SLs in either location strongly inhibited 48S PIC formation. Ded1 increased the maximal rate of *CP-8.1* recruitment by ~5-fold, such that the maximal rate for *CP-8.1* was still ~6-fold below that of *-SL* (Figs. 2C; S2F, blue vs. red), and ~2-3-fold below that of *RPL41A, HOR7, SFT2, OST3*, and *PMA1* (Figs. 2C, 1B). Moreover, *CP-8.1* exhibited an ~7.6-fold higher 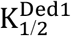 versus *-SL* (Fig. 2D). As a result, *CP-8.1* lies between the Ded1-hyperdependent and Ded1-hypodependent mRNAs, and deviates from the line in the plot of 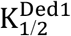 versus fold-increase in k_max_ (Fig. 2E, green, *CP-8.1*). This deviation is due to the fact that *CP-8.1*, like the Ded1-hyperdependent mRNAs, is inefficiently recruited in the absence of Ded1; but unlike the Ded1-hyperdependent mRNAs, its recruitment is accelerated by Ded1 only ~5-fold. These results suggest that Ded1 is not very effective at reducing the inhibitory effect of a stable cap-proximal SL on PIC attachment, even at saturating Ded1 concentrations. In contrast, Ded1 conferred an ~17-fold acceleration of recruitment for *CD-8.1* (Fig. 2C), and a high Ded1 concentration was required to achieve the half-maximal rate (Figs. 2D; S2F, green vs. red), similar to the behavior of the native Ded1-hyperdependent mRNAs (Figs. 1B-C). Judging by the relative positions of *CD-8.1* and *CP-8.1* on the plot of Fig. 2E, it appears that Ded1 is better at resolving the inhibitory effect of the synthetic SL in a cap-distal versus cap-proximal location in the 5′-UTR. Interestingly, the same conclusion was reached previously from analyzing the relative effects of a cold-sensitive *ded1* mutation on expression of reporter mRNAs containing cap-proximal or cap-distal SLs (Sen et al., 2015).

Having observed that the strength of a 5′-UTR SL influences the degree of Ded1-dependence, as observed with WT versus mutant derivatives of *SFT2* and *PMA1* (Fig. 2C), we went on to analyze synthetic mRNAs containing cap-distal SLs of predicted stabilities either higher (−10.5 kcal/mol) or lower (−3.7 kcal/mol) than that of *CD-8.1.* Each of these SLs, present in *CD-10.5* and *CD-3.7*, respectively, reduced the recruitment rate in the absence of Ded1 by ~20-fold compared to that of *-SL*, similar to the results for *CD-8.1* (Fig. 2F, blue). This result suggests that eIF4E·eIF4G·eIF4A alone cannot efficiently mitigate the inhibitory effects of cap-distal SLs of even moderate stability, such as that in *CD-3.7*, consistent with previous findings (Yourik et al., 2017). As expected, Ded1 strongly stimulated the recruitment rate of *CD-3.7*, by ~25-fold, slightly more than the ~15-fold observed for *CD-8.1* (Fig. 2F); however, Ded1 conferred only a modest ~2-fold acceleration of *CD-10.5* recruitment (Fig. 2F). One possibility to explain these last results would be that the cap-distal SL in *CD-10.5* is too stable for efficient unwinding by Ded1, limiting Ded1’s ability to accelerate 48S assembly on this mRNA.

In summary, our analysis of synthetic mRNAs supports the notion that Ded1 can accelerate mRNA recruitment by enhancing scanning through cap-distal secondary structures, such as the SLs that occur naturally in *SFT2* and *PMA1* mRNAs; although if the structure is too stable, Ded1 has a limited ability to unwind it. Additionally, it appears that Ded1 also partially mitigates the inhibitory effects of cap-proximal secondary structures on PIC attachment at the mRNA 5′-end, although not to the same degree that it overcomes cap-distal structures.

### Ded1 stimulation is completely dependent on eIF4G·eIF4E for a subset of mRNAs and all mRNAs require eIF4A in the presence or absence of Ded1

Ded1 has been shown to interact physically with eIF4F components eIF4G and eIF4A in a manner that influences its ability to unwind model RNA substrates (Gao et al., 2016; Hilliker et al., 2011; Senissar et al., 2014). Accordingly, we examined whether its interactions with eIF4F influence Ded1’s ability to accelerate 48S PIC assembly. As eIF4E is co-purified with eIF4G (Mitchell et al., 2010), and eIF4E is required for full activity of the eIF4F complex, all experiments involving eIF4G utilize the eIF4E·eIF4G heterodimer, hereafter referred to simply as eIF4G. We performed mRNA recruitment assays in the presence and absence of eIF4G and Ded1 for (i) Ded1-hypodependent mRNAs *RPL41A, HOR7, -SL*, and *SFT2-M;* (ii) Ded1-hyperdependent mRNAs *SFT2, OST3*, and *CD-8.1;* and (iii) *CP-8.1*, which exhibits intermediate behavior between groups (i) and (ii) mRNAs in the relationship between 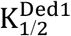 and k_max_ stimulation (Fig. 2E). As described above, in the presence of eIF4G, all eight mRNAs can be recruited in the absence of Ded1, and Ded1 increased their recruitment rates to different extents (Figs. 3A-H, blue vs. orange). *RPL41A, HOR7*, and *-SL*, which contain very short (*RPL41A* and *HOR7*) or unstructured (*-SL*) 5′-UTRs, were recruited at relatively low (but measurable) rates in the absence of both eIF4G and Ded1 (Figs. 3A, 3B, 3F, tan bars); and Ded1 conferred no increase in their apparent rates in the absence of eIF4G (Figs. 3A, 3B, 3F, cf. green vs. tan, orange vs. blue). The complete dependence of Ded1 on eIF4G to accelerate recruitment for these three mRNAs is consistent with the idea that Ded1 acts exclusively in the context of the eIF4GeIF4EeIF4ADed1 quaternary complex (Gao et al., 2016). The same might be true for *CP-8.1*, whose observable stimulation by Ded1 also required eIF4G (Fig. 3H); however, because no *CP-8.1* recruitment was observed without eIF4G, Ded1 might stimulate recruitment of this mRNA on its own at levels below the detection limit of the assay. The finding that *CP-8.1* differs from *-SL* in showing no measurable recruitment in the absence of eIF4G (Figs. 3H vs 3F), suggests that the cap-proximal SL in *CP-8.1* imposes a requirement for eIF4G.

**Fig. 3:**
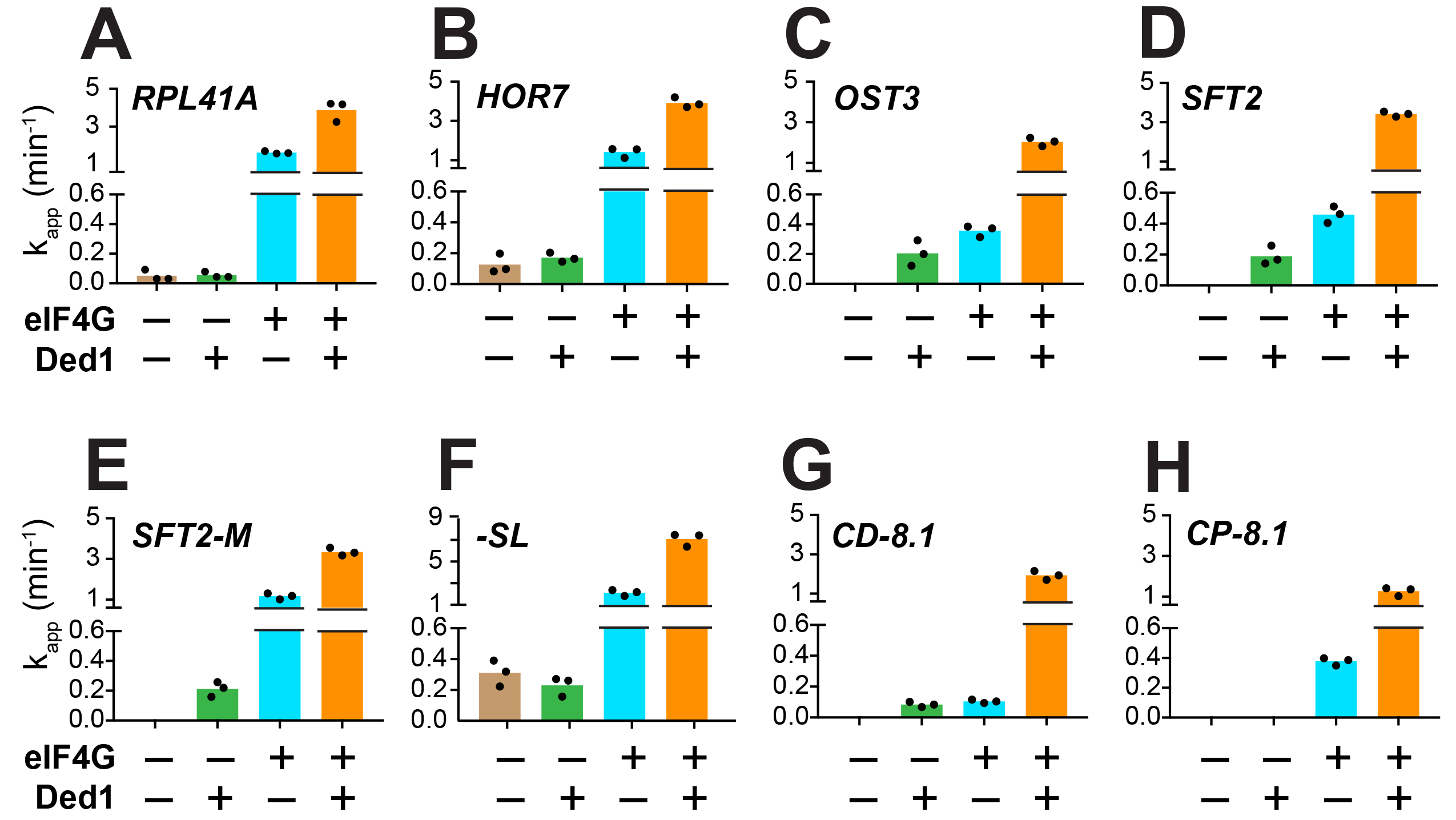
Ded1 acceleration of 48S PIC assembly is completely dependent on eIF4G for a subset of mRNAs. **(A-H)** Apparent rates (k_app_) of 48S PIC assembly for the indicated mRNAs observed without eIF4G and Ded1 (tan bars), with Ded1 but no eIF4G (green bars), with eIF4G but no Ded1 (blue bars), or with both Ded1 and eIF4G (orange bars). Note the different Y-axis scale for *-SL* (F). See Supplementary Figure 3 and Figure3_SourceData1

In contrast to the results described above, Ded1 can accelerate recruitment of *SFT2, OST3, SFT2-M*, and *CD-8.* l mRNAs in the absence of eIF4G, with comparable apparent rates afforded by eIF4G alone for *SFT2, OST3*, and *CD-8.* l (Figs. 3C, 3D, 3G; green vs. blue), but well below the apparent rates observed with eIF4G for *SFT2-M* (Fig. 3E, green vs. blue). Thus, Ded1 can stimulate recruitment of these four mRNAs, at least to some extent, acting outside of the eIF4G·eIF4E·eIF4A·Ded1 complex.

In contrast to eIF4G, omitting eIF4A from the reactions essentially eliminated recruitment of all mRNAs tested, yielding endpoints of <10%; with the exception of *-SL*, which was recruited at the low rate of 0.04 ± 0.01 min^−1^ with endpoints of 46 ± 3% (Fig. S3, red). Moreover, in the absence of eIF4A, Ded1 did not rescue recruitment of any mRNAs, nor did it increase the k_app_ or endpoints for *-SL* mRNA (Fig. S3, blue). Thus, eIF4A has one or more essential functions in mRNA recruitment that cannot be provided by Ded1, even for an mRNA such as *-SL* exhibiting appreciable recruitment in the absence of eIF4G (Fig. 3F, tan). This is consistent with the previous findings that eIF4A is required for robust translation of virtually all yeast mRNAs in vivo and in vitro (Sen et al., 2015; Yourik et al., 2017), and that DHX29 and yeast Ded1 cannot substitute for eIF4A in 48S PIC assembly on native β-globin mRNA in a mammalian reconstituted system (Abaeva et al., 2011; Pisareva et al., 2008).

### Interaction between the RNA3 domain of eIF4G and Ded1-CTD stimulates mRNA recruitment

The C-terminal RNA3 domain of eIF4G was shown to interact physically with the Ded1 CTD (Hilliker et al., 2011) and to influence effects of eIF4G on Ded1 unwinding of a model RNA duplex in vitro (Gao et al., 2016; Putnam et al., 2015). Hence, we sought to determine whether this physical interaction between the CTDs of Ded1 and eIF4G is functionally relevant in 48S PIC assembly by performing mRNA recruitment assays with a truncated eIF4G variant lacking RNA3 (eIF4G·ΔRNA3) or a Ded1 variant lacking the CTD (Ded1Δ-CTD) (Fig. S4A).

Without Ded1, the ΔRNA3 truncation of eIF4G had little or no effect on k_max_ for all seven mRNAs tested (Fig. 4A, compare blue bars to superimposed line/whiskers, the latter indicating results for WT eIF4G re-plotted from Fig. 3A-H for comparison, where the horizontal line shows the mean and the whiskers one SD from the mean). In the presence of Ded1, by contrast, ΔRNA3 conferred ~2-fold reductions in k_max_ for three mRNAs, *RPL41A* (ΔRNA3 - 1.5 ± 0.2 min^−1^; WT - 3.9 ± 0.5 min^−1^), *HOR7* (ΔRNA3 - 1.8 ± 0.1 min^−1^ WT - 3.9 ± 0.3 min^−1^) and *CP-8.1* (ΔRNA3 - 0.7 ± 0.03 min^−1^; WT - 1.3 ± 0.1 min^−1^), compared to the values observed for WT eIF4G (Fig. 4A, orange bars vs. line/whiskers). Somewhat smaller reductions in k_max_ were observed for *SFT2* and *SFT2-M* (Fig. 4A). Thus, as summarized in Fig. 4B, deleting RNA3 nearly eliminated the stimulatory effect of Ded1 on k_max_ values for *RPL41A, HOR7*, and *CP-8.1* (black vs. grey bars vs. dashed red line, the latter indicating no stimulation by Ded1).

**Fig. 4:**
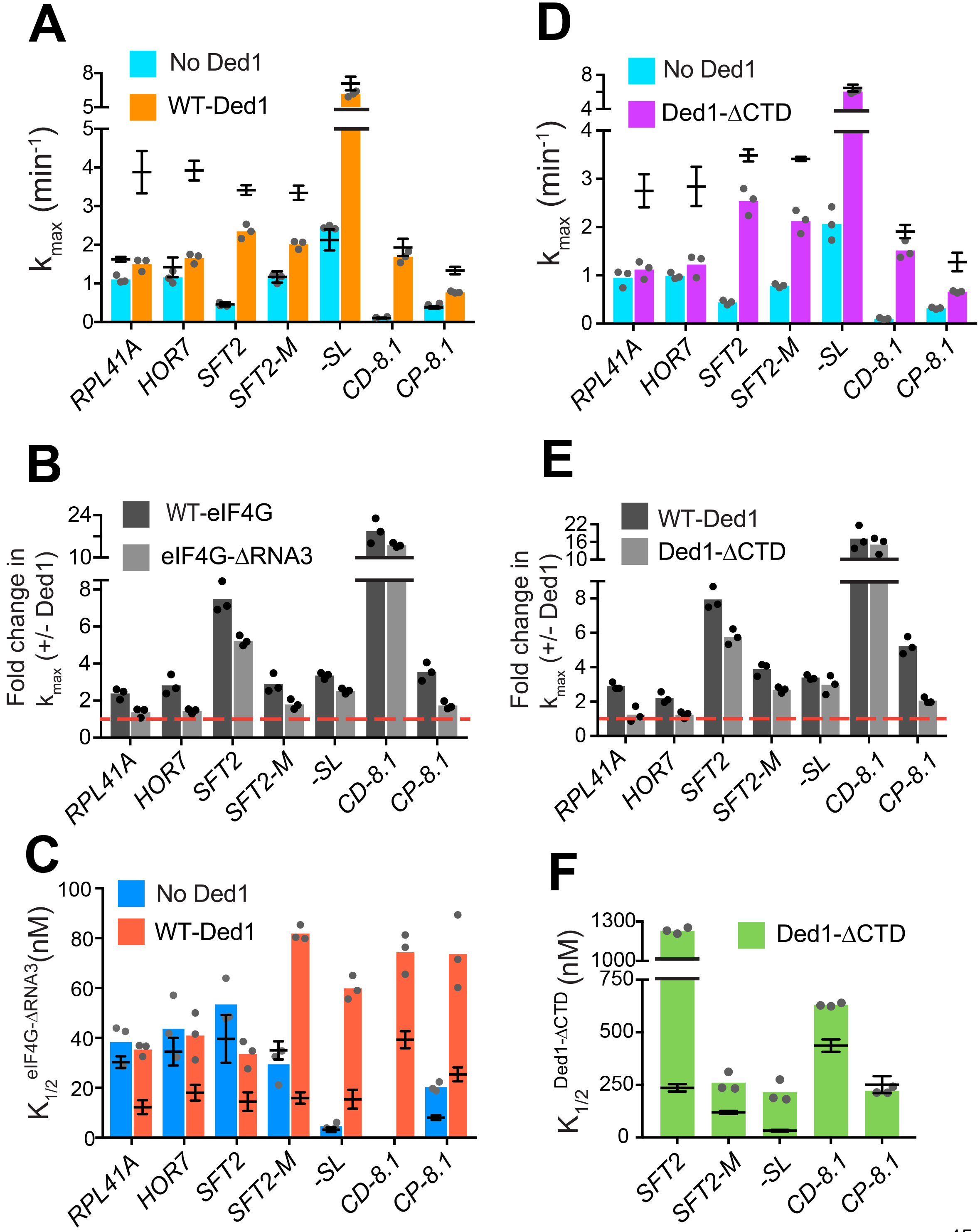
eIF4G-RNA3 and Ded1-CTD enhance k_max_ for the same subset of mRNAs while reducing K1/2 for nearly all mRNAs. **(A)** k_max_ values in absence (blue) and presence of saturating Ded1 (orange) with eIF4G-ΔRNA3. **(B)** The average fold-change in the k_max_ observed in the presence and absence of Ded1 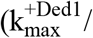 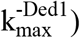 with WT eIF4G (dark grey) and eIF4G-ΔRNA3 (light grey). Red line indicates no change in the rate on Ded1 addition. **(C)** K_1/2_ of eIF4G-ΔRNA3 in absence of Ded1 (dark blue) and presence of saturating Ded1 (red). **(D)** k_max_ observed in absence (blue) and presence of saturating Ded1-ΔCTD (purple). **(E)** The average fold change in the k_max_ 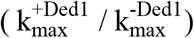 observed in presence and absence of WT Ded1 (dark grey) and Ded1-μCTD (light grey). Red line indicates rates in absence of Ded1. **(F)** The K_1/2_ of Ded1-ΔCTD value shown as green bars. (A-F) Bars indicate mean values calculated from the 3 independent experiments represented by the data points. The superimposed horizontal line (black) indicate the mean maximal rates (A, D) or K_1/2_ (C, F) observed with WT eIF4G (A, C, from Figs. 3 and S5A) or WT Ded1 (D, F from Figs. 1 and 2), and error bars represent 1 SD from the mean (this representation will be referred to as line/whisker plot). See Supplementary Figures 4 and 5 and Figure4_SourceData1

Importantly, when Ded1 was replaced with Ded1-ΔCTD in reactions containing WT eIF4G, we observed effects on k_max_ values and rate enhancements very similar to those described above for ΔRNA3 (Figs. 4D, 4E), which is consistent with a functional interaction between RNA3 and Ded1-CTD. (We verified that saturating concentrations of eIF4G-ΔRNA3 and Ded1-ΔCTD were used in these experiments by showing that rates of recruitment for *SFT2, RPL41A* or *HOR7* mRNAs were not elevated even at much higher concentrations of the variants (Figs. S4B, S4C, S4D, and data not shown). We also confirmed that Ded1-ΔCTD has RNA-dependent ATPase activity similar to that of the WT (Figs. S1B-C)).

We next examined whether eliminating eIF4G-RNA3 alters the concentration of eIF4G required to achieve half-maximal rate acceleration, ie. the 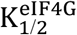. To this end, we first determined the 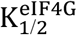 of WT eIF4G for each mRNA in the presence or absence of Ded1. For four native mRNAs and *SFT2-M*, the presence of Ded1 lowered the 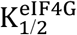 by factors of ~2-to 2.5 (Fig. S5A, cols. 1-5, red vs. blue), consistent with the idea that Ded1 interacts productively with eIF4G on all five of these mRNAs. Contrary to the natural mRNAs, addition of Ded1 increased the 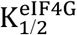 of *-SL* and *CP-8.1* mRNAs, reaching values similar to those observed for the natural mRNAs in the presence of Ded1 (Fig. S5A, cols. 6 & 8, red vs. blue). (The 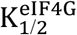 for *CD-8.1* without Ded1 could not be determined accurately because of its endpoint defects at lower eIF4G concentrations.) Thus, the maximum stimulation of recruitment of the synthetic mRNAs in the absence of Ded1 can be achieved at relatively low eIF4G concentrations, but higher eIF4G concentrations are required to support the additional stimulation of recruitment conferred by Ded1. (See Fig. S5 legend for additional comments.)

We then proceeded to determine the effect of eliminating RNA3 on the concentration of eIF4G required for the maximum recruitment rate in the absence of Ded1. For mRNAs *RPL41A, HOR7, SFT2, SFT2-M*, and *-SL*, the K_1/2_ values for eIF4G-ΔRNA3 did not differ substantially from those of WT eIF4G, although it was ~2-fold higher for *CP-8.1* (Fig. 4C, blue bars vs. line/whiskers summarizing results for WT eIF4G taken from Fig. S5A). (The 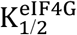 could not be accurately measured for *CD-8.1* using eIF4G-ΔRNA3 due to endpoint defects.) In reactions containing Ded1, by contrast, the 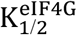 values for eIF4G-ΔRNA3 were increased by 2- to 5fold relative to the values determined for WT eIF4G for all mRNAs tested (Fig. 4C, red bars versus superimposed line/whiskers results for WT eIF4G taken from Fig. S5A). Thus, on removal of RNA3, relatively higher concentrations of eIF4G are required to achieve maximal rate stimulation by Ded1, supporting a functionally important interaction between RNA3 and Ded1. We also determined the effects of eliminating the Ded1 CTD on K_1/2_ values for Ded1. Similar to the results obtained for eIF4G-ΔRNA3, higher K_1/2_ values were observed for Ded1-ΔCTD versus WT Ded1 for four of the five mRNAs that exhibit appreciable rate stimulation by Ded1-ΔCTD (which excludes *RPL41A* and *HOR7*) (Fig. 4F, green bars versus superimposed line/whiskers results for WT Ded1 from Figs. 1C & 2D).

In summary, our results indicate that there is an interaction between eIF4G-RNA3 and Ded1-CTD that facilitates Ded1 function in mRNA recruitment to the PIC. The increased K_1/2_ values evoked by eliminating either domain suggests that their interaction enhances assembly of the eIF4G·eIF4E·eIF4A·Ded1 tetrameric complex (Gao et al., 2016). The finding that increased concentrations of eIF4G-ΔRNA3 or Ded1-ΔCTD can rescue the rate (k_max_) defects caused by the domain deletions for some mRNAs but not for others suggests that in certain mRNA contexts this interaction plays a role in addition to simply promoting interaction between eIF4G and Ded1.

### The RNA2 domain of eIF4G functions in Ded1-dependent mRNA recruitment

Ded1 also interacts with the RNA2 domain of eIF4G (Senissar et al., 2014), but the importance of this interaction for Ded1 function is unknown. Comparing the eIF4G-ΔRNA2 variant (with an internal deletion of RNA2) to WT eIF4G, we found that, as for RNA3, deletion of RNA2 influenced recruitment of the mRNAs to different extents. In reactions lacking Ded1, we observed no significant differences in k_max_ for any of the seven mRNAs examined (Fig. 5A, blue bars vs. line/whiskers for WT eIF4G data from Fig. 3, blue). By contrast, removal of RNA2 increased the K_1/2_ for eIF4G-ΔRNA2 versus WT eIF4G by 3-5-fold for *SFT2-M, -SL*, and *CP-8.1* mRNAs in reactions lacking Ded1 (Fig. 5C, blue bars vs. line/whiskers from Fig. S5A). ΔRNA2 also conferred an endpoint defect for *SFT2* at lower concentrations, precluding determination of its effects on the K_1/2_ for eIF4G. (Because of endpoint defects for *CD-8.1* even with WT eIF4G, the importance of RNA2 cannot be evaluated for this mRNA.) Considering that ΔRNA3 increased the K_1/2_ for eIF4G only for *CP-8.1* in reactions lacking Ded1, it appears that RNA2 has relatively more important Ded1-independent functions than RNA3 in recruitment of particular mRNAs.

**Fig. 5:**
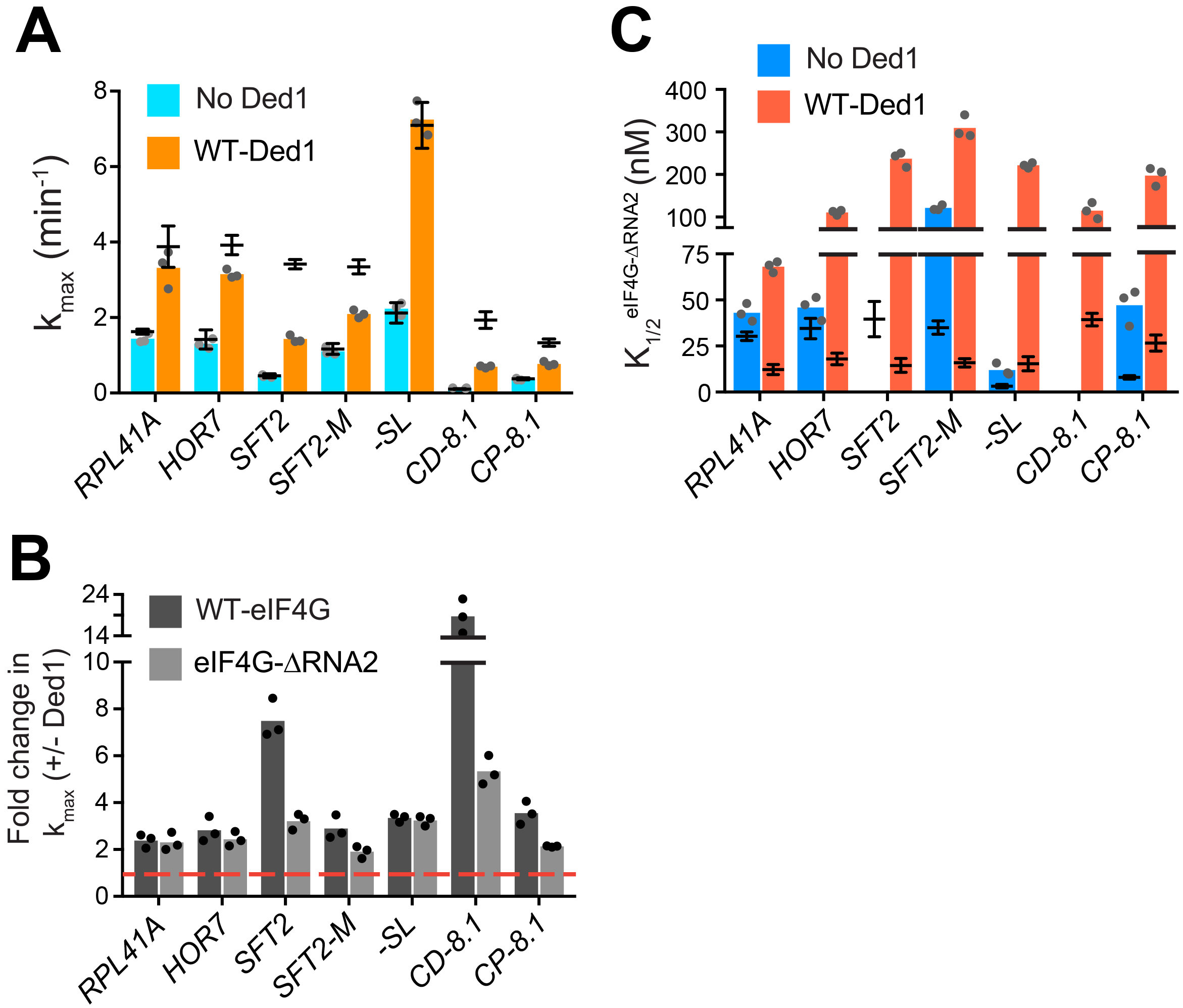
RNA2 domain of eIF4G is crucial for Ded1-dependent stimulation of mRNA recruitment. **(A)** k_max_ in the absence (blue) and presence of saturating Ded1 (orange) with eIF4G-ΔRNA2. **(B)** The average fold-change in the maximal rate observed in the presence and absence of Ded1 with WT eIF4G (dark grey) and eIF4G-ΔRNA2 (light grey). Red line indicates no change in the rate on Ded1 addition. **(C)** The K_1/2_ of eIF4G-ΔRNA2 in the absence of Ded1 (dark blue) and presence of saturating Ded1 (red). (A-C) Bars represent mean values (n = 3) and points on each bar show the individual experimental values. The line and whisker plot indicate the mean maximal rates (A) or K_1/2_ (C) observed with WT eIF4G, and error bars represent 1 SD, as depicted in Fig. 4. See Supplementary Figures 4 and 5 and Figure5_SourceData1

Stronger effects of μRNA2 were observed in reactions containing Ded1, markedly reducing the k_max_ for the two mRNAs containing cap-distal SLs, *SFT2* (k_max_ of 1.4 ± 0.1 min^−1^ vs. 3.4 ± 0.1 min^−1^ for WT) and *CD-8.1* (k_max_ of 0.7 ± 0.03 min^−1^ vs. 1.9 ± 0.2 min^−1^ for WT), with smaller reductions for *SFT2-M* and *CP-8.1* (Fig. 5A, orange bars vs. line/whiskers; Fig. 5B, black vs. grey bars vs. dotted red line). By contrast, RNA2 was dispensable for maximal Ded1 acceleration of *RPL41A, HOR7*, and *-SL* mRNA recruitment (Fig. 5B, black vs. grey bars). It is noteworthy that *RPL41A* and *HOR7* exhibited the strongest dependence on RNA3 (Fig. 4B), but were insensitive to loss of RNA2 (Fig. 5B) for maximal rate stimulation by Ded1.

All of the mRNAs exhibited increases in K_1/2_ for eIF4G-ΔRNA2 versus WT eIF4G (320-fold; Fig. 5C, red bars vs. line/whiskers). Thus, eliminating RNA2 increases the concentration of eIF4G required for maximal Ded1 stimulation for all mRNAs tested, which might reflect its importance in promoting formation of the eIF4G·eIF4E·eIF4A·Ded1 complex, in the manner concluded above for RNA3. Elevated concentrations of eIF4G-ΔRNA2 enable maximum recruitment rates similar to those achieved with WT eIF4G for the mRNAs with lower degrees of structure - *RPL41A, HOR7*, and *-SL;* whereas the more structured mRNAs display varying reductions in k_max_ at saturating eIF4G-ΔRNA2 concentrations, with the two mRNAs harboring cap-distal stem loops -*SFT2* and *CD-8.1* - having the largest rate enhancement defects. This suggests that RNA2 enhances Ded1 function on the structured mRNAs beyond its ability to simply stabilize Ded1-eIF4G interaction.

### The N-terminal domain of Ded1 enhances mRNA recruitment

It was shown previously that the N-terminal domain (NTD) of Ded1 physically interacts with eIF4A, and is required for eIF4A stimulation of Ded1 unwinding activity in vitro (Gao et al., 2016; Senissar et al., 2014). We tested the effects of Ded1 on K_1/2_ of eIF4A and, as observed with WT eIF4G, Ded1 influenced 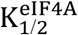 differently on these mRNAs, providing evidence that Ded1 has functional interactions with eIF4A during mRNA recruitment (Fig. S5B). Hence, we examined the effect of eliminating the Ded1-NTD on recruitment of our panel of mRNAs. The k_cat_ and K_m_ values for RNA-dependent ATP hydrolysis were indistinguishable between WT Ded1 and the ΔNTD variant (Figs. S1C-D). For the *SFT2, CD-8.1*, and *CP-8.1* mRNAs, which harbor defined SLs in their 5′-UTRs, eliminating the Ded1 NTD decreased k_max_ by l.5-2-fold, whereas the k_max_ values for *RPL41A, HOR7, SFT2-M*, and *-SL* mRNAs were not significantly altered by ΔNTD (Figs. 6A, purple bars vs. line/whiskers for WT Ded1; Fig. 6B). However, 1-2 orders of magnitude higher K_1/2_ values for the Ded1-ΔNTD versus WT Ded1 were observed with all mRNAs except *CD-8.1* (Fig. 6C, green bars vs. line/whiskers for WT Ded1). Thus, removing the Ded1 NTD significantly increases the concentrations of Ded1 required to achieve enhancement of recruitment of six out of the seven mRNAs tested. These data are consistent with the idea that interaction of eIF4A and the Ded1 NTD enhances assembly or stability of the eIF4GΔeIF4EΔeIF4AΔDed1 complex, and stimulates Ded1 helicase activity; although the Ded1-NTD might also mediate important interactions with other components of the system. As with deletions of the eIF4G RNA domains and Ded1-CTD, mRNA-specific defects were conferred by deleting the Ded1 NTD.

**Fig. 6:**
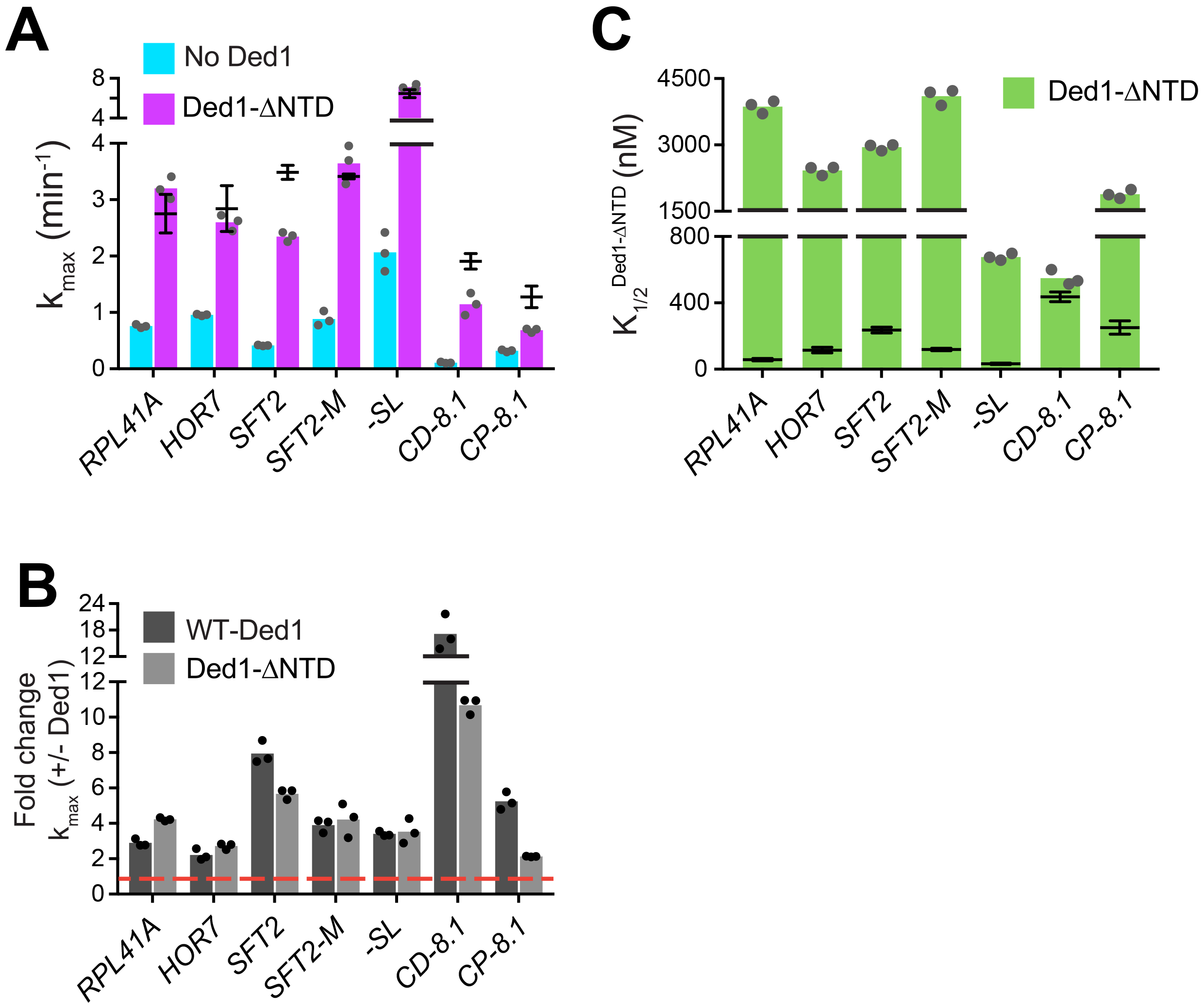
Ded1-NTD enhances mRNA recruitment. **(A)** k_max_ values observed in the absence (blue) and presence of saturating Ded1-ΔNTD (purple). **(B)** The average fold change in the maximal rate 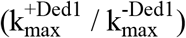 observed in the presence and absence of WT Ded1 (dark grey) and Ded1-ΔNTD (light grey). Red line indicates no change in the rate with Ded1. **(C)** K_1/2_ of Ded1-ΔNTD value shown as green bars. (A-C) Bars represent the mean values (n = 3) and points on each bar show the individual experimental values. The line and whisker plot indicate the mean maximal rates (A) or K_1/2_ (C) observed with WT Ded1, and error bars represent 1 SD. See Supplementary Figures 4 and 5 and Figure6_SourceData1

## DISCUSSION

Employing a purified yeast translation initiation system, we reconstituted the function of DEAD-box helicase Ded1 in stimulating the rate of 48S PIC assembly on both native and model mRNAs. This stimulation in vitro recapitulates the Ded1-dependence of translation of mRNAs observed in vivo using ribosome profiling, in which mRNAs having longer and more structured 5′-UTRs display hyper-dependence on Ded1 relative to mRNAs with shorter and less structured 5′-UTRs (Sen et al., 2015). We showed that defined SL structures both decrease rates of 48S PIC assembly in the absence of Ded1 and increase the fold-stimulation afforded by Ded1. These results provide direct biochemical evidence supporting the proposition that Ded1 enhances translation initiation in vivo by resolving secondary structures formed by 5′-UTR sequences. Our results also showed that Ded1-accelerated recruitment of several mRNAs depends completely on the presence of eIF4G, and that domains mediating Ded1 interactions with eIF4G or eIF4A enhance Ded1 stimulation of 48S PIC assembly, consistent with the conclusion that Ded1 binding to the eIF4G·eIF4E·eIF4A complex enhances its activity (Gao et al., 2016). However, Ded1 can also stimulate recruitment of some mRNAs in the absence of eIF4G, indicating that it can act independently of eIF4F as well.

### DEAD-box proteins Ded1 and eIF4A have complementary but distinct functions in mRNA recruitment

Inactivation of conditional Ded1 mutants in vivo results in a strong reduction in bulk polysomes and decreased expression of reporter mRNAs bearing unstructured 5′-UTRs (Chuang et al., 1997; de la Cruz et al., 1997; Sen et al., 2015), suggesting that Ded1 promotes translation of most mRNAs, in addition to playing a more specific role for mRNAs with long, structured 5′-UTRs. Consistent with this, we observed that Ded1 increases the maximal rate of 48S PIC formation by ~2-3-fold on mRNAs with 5′-UTRs with low degrees of secondary structure (*RPL41A, HOR7*, and *SFT2-M)*, as well as on a synthetic mRNA with an unstructured 5′-UTR (*-SL*). eIF4A also enhances the translation of nearly all mRNAs in vivo, although, unlike Ded1 where sizable sets of mRNAs are hyper- or hypo-dependent on its function, most mRNAs are similarly (strongly) dependent on eIF4A for translation (Sen et al., 2015). In line with these observations, in vitro, eIF4A promotes 48S PIC assembly on all mRNAs tested, increasing the k_max_ for the synthetic mRNA with unstructured 5′-UTR (*-SL*) by 60-fold and even accelerating recruitment of completely unstructured model mRNAs by ≥ 7-fold (Yourik et al., 2017). However, although Ded1 and eIF4A both facilitate recruitment of most mRNAs, their functions are distinct. Both proteins are essential in vivo (Chuang et al., 1997; Linder and Slonimski, 1989); and Ded1 cannot substitute for eIF4A in vitro (Fig. S3), but it promoted recruitment of all mRNAs tested beyond the level achieved by saturating concentrations of eIF4A and eIF4G·eIF4E (Figs. 1B, 2C).

We previously proposed that eIF4A stimulates a step of 48S PIC assembly common to all mRNAs, such as disrupting the ensemble of weak RNA-RNA interactions that impede PIC attachment to the 5′-UTR or subsequent scanning (Yourik et al., 2017). In addition, eIF4A might also directly promote loading of mRNA onto the PIC, e.g. by modulating conformational changes in the 40S subunit or by threading the 5′-end into the mRNA binding channel (Kumar et al., 2016; Sokabe and Fraser, 2017). In common with eIF4A, Ded1 may enhance recruitment of all mRNAs by disrupting their global structures created by dynamic ensembles of base-pairing throughout their lengths. Unlike eIF4A, however, Ded1 can efficiently resolve more stable structures, including local stem-loops—achieving an order-of-magnitude acceleration for mRNAs with the most structured 5′-UTRs. If the proposed Ded1 function in promoting 48S PIC formation by disrupting global mRNA structure requires lower Ded1 concentrations than its role in resolving more stable structures within or involving the 5′-UTR, it would be consistent with our findings that mRNAs with SLs require higher Ded1 concentrations to achieve the much greater fold-stimulation of 48S assembly afforded by Ded1 compared to mRNAs lacking SLs (Figs. 1-2).

### Evidence supporting the functional importance of an eIF4G·eIF4E·eIF4A-Ded1 tetrameric complex

Ded1 alters the K_1/2_ of eIF4G and eIF4A for most mRNAs (Fig. S5), and these new K_1/2_ values may signify the changes in the concentrations of eIF4A and eIF4G required for proper assembly of the eIF4G·eIF4E·eIF4A·Ded1 tetrameric complex on each mRNA. The effects of eliminating known interactions between Ded1 and eIF4G or eIF4A further suggests the importance of the tetrameric complex formation for Ded1 function. With only two exceptions (*CP-8.1* for Ded1-ΔCTD and *CD-8.1* for the Ded1-ΔNTD), we found that deleting the RNA2 or RNA3 domain of eIF4G, or the CTD or NTD of Ded1, increased the concentrations of the corresponding eIF4G or Ded1 variants required to achieve the half-maximal rate of 48S PIC assembly (ie., their K_1/2_ values) on each mRNA examined, as summarized by the heatmap in Fig. 7A. This holds for the mRNAs with the shortest or least structured 5′-UTRs (*-SL, RPL41A*, and *HOR7*) as well as those with the most highly structured 5′-UTRs (*SFT2, CD-8.1*, and *CP-8.1*). Because all of these domain deletions abrogate known interactions linking Ded1 to eIF4G or eIF4A, a plausible way to account for these findings is to propose that, regardless of the amount of secondary structure in the mRNA 5′-UTR, rapid mRNA recruitment depends on Ded1 functioning within the eIF4E·eIF4G·eIF4A·Ded1 complex; and that eliminating any interaction between Ded1 and eIF4G or eIF4A necessitates a higher concentration of the mutant variant for efficient complex formation. Judging by the magnitude of the increases in K1/2 conferred by eliminating different domains (Fig. 7A), it would appear that eIF4G-RNA2 and Ded1-NTD are generally more important than the eIF4G-RNA3/Ded1-CTD duo in promoting assembly or stability of the eIF4E·eIF4G·eIF4A·Ded1 complex. However, we cannot rule out the possibility that the domain deletions also impair a different interaction, for instance, with mRNA or another factor such as eIF4B or eIF3 that is crucial for rapid mRNA recruitment (Mitchell et al., 2010).

**Fig. 7:**
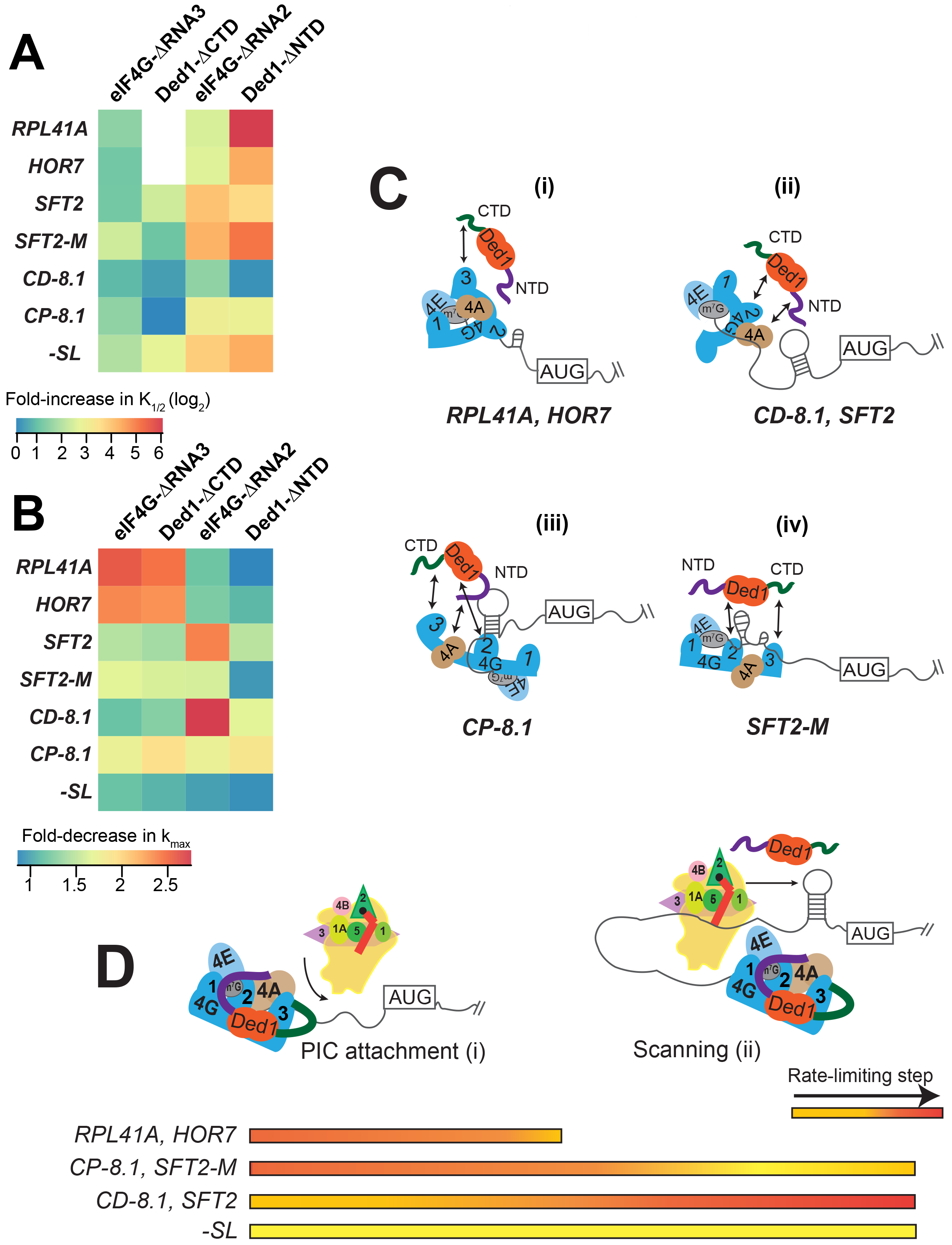
Models for mRNA-specific eIF4F-Ded1 interactions. **(A)** Heatmap representation of log_2_ fold-increases in K_1/2_ values of eIF4G-ΔRNA3 or eIF4G-ΔRNA2 versus WT eIF4G; and of Ded1-ΔCTD or Ded1-ΔNTD versus WT Ded1, calculated from data in Figs. 4C & 5C (red bars vs. line/whiskers) or Figs. 4F & 6C (green bars vs. line/whiskers). **(B)** Heatmap depicting the fold-changes in k_max_ values of the indicated eIF4G or Ded1 truncations 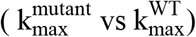 summarized calculated from Figs. 4A, 4D, 5A, and 6A (orange or purple bars vs line/whiskers). **(C)** mRNA “geometry” model depicting how different mRNAs exhibit distinct configurations regarding the occurrence and location of RNA structures (shown as hairpins or stem-loops) that influence the relative importance of different domain interactions linking Ded1 to eIF4G or eIF4A within the eIF4G·eIF4E·eIF4A·Ded1 tetrameric complex. **(D)**“Kinetic” model depicting how different mRNAs differ in the extent to which PIC attachment or scanning are rate-limiting steps in 48S PIC assembly, which can be modulated by Ded1 acting alone or within the eIF4E eIF4G·eIF4E·eIF4A·Ded1 complex.

### Ded1 and eIF4G mutants affect the maximal rates for recruitment differently depending on the mRNA

With some mRNAs, the maximum rate of recruitment observed with WT Ded1 and WT eIF4G (k_max_) could also be achieved using elevated concentrations of the mutant variant; whereas in other cases, the observed k_max_ was diminished from the WT value even at saturating amounts of the mutant Ded1 or eIF4G variant. We regard such reductions in k_max_ as indicating impairment of a fundamental role of the deleted eIF4G or Ded1 domain in rapid recruitment of the affected mRNA. Accordingly, we summarized these effects for each factor truncation in a heatmap (Fig. 7B) to evaluate the requirements for particular domains or interactions for maximum rate stimulation by Ded1 with each mRNA tested.

It is evident from the heatmap that deletion of the eIF4G-RNA3 or Ded1-CTD domain confers a wide range of k_max_ reductions: 2-3-fold for *RPL41A* and *HOR7*; ~1.5-fold for *SFT2, SFT2-M*, and *CP-8.1*, and almost no change for *-SL* and *CD-8.1* (Fig. 7B, cf. cols. 1-2, all rows). Importantly, however, in all cases the effects of deleting eIF4G-RNA3 and Ded1-CTD are similar for a given mRNA, supporting the proposition that these effects result from loss of the eIF4G-RNA3/Ded1-CTD interaction. Eliminating this interaction essentially abolishes the rate enhancement provided by Ded1 for recruitment of *RPL41A* and *HOR7* mRNAs (Figs. 4B, 4E), indicating that the Ded1-CTD/eIF4G-RNA3 interaction is essential for the eIF4E·eIF4G·eIF4A·Ded1 complex (i.e., at saturating Ded1 concentration) to accelerate a slow step in 48S PIC assembly on these mRNAs (Fig. 7C(i)). This step is apparently enhanced through different interactions or is less rate-limiting for the other mRNAs tested.

We observed a similar diversity of effects on k_max_ values depending on the mRNA for the eIF4G-RNA2 and Ded1-NTD deletions. ΔRNA2 had little effect on k_max_ for *RPL41A* and *HOR7*, in contrast to the deleterious effects of ΔRNA3 for these two mRNAs. Similarly, ΔRNA2 markedly reduced the k_max_ for *SFT2* and *CD-8.1* by ~2.5-3-fold, whereas ΔRNA3 had little effect on these mRNAs (Fig. 7B, col. 3 vs. col. 1). Similar to ΔRNA3 however, ΔRNA2 decreased the k_max_ values for *SFT2-M* and *CP-8.1* by ~1.5-2-fold, with minimal effect on *-SL* mRNA (Fig. 7B). In this case, the RNA2 domain appears to be most important for the ability of the eIF4E·eIF4G·eIF4A·Ded1 complex to stimulate recruitment of the two mRNAs with cap-distal SLs, *SFT2* and *CD-8.1*; nearly dispensable for the mRNAs with the lowest degrees of structure, - *SL, RPL41A* and *HOR7;* and of intermediate importance for *CP-8.1* and *SFT2-M.* These observations suggest that RNA2 facilitates Ded1 function in melting out structures encountered by the PIC during attachment or scanning (Fig. 7C (ii-iv)). (Although *SFT2-M*lacks the major cap-distal SL in WT *SFT2*, it contains an additional cap-proximal structure in vitro that might underlie its greater dependence on RNA2 versus *-SL, RPL41A*, and *HOR7.*)

Whereas deletion of the Ded1-NTD had the largest effect of the four eIF4G or Ded1 truncations on K_1/2_ values (Fig. 7A), it had the smallest effects on maximal rates of recruitment, reducing k_max_ values between ~1.5 to ~2-fold for *SFT2, CD-8.1*, and *CP-8.1*, but having little effect on the other mRNAs (Fig. 7B). As all three affected mRNAs have stable SLs and the unaffected mRNAs do not, these data might indicate that the Ded1-NTD, presumably by interacting with eIF4A, modestly enhances the ability of the eIF4E·eIF4G·eIF4A·Ded1 complex to unwind stable secondary structures—complementing the function of eIF4G-RNA2 in this reaction (Fig. 7C (ii-iii)).

What is perhaps most striking about the effects of the eIF4G and Ded1 truncations on both the K_1/2_ (Fig. 7A) and k_max_ (Fig. 7B) values is that mRNAs have distinct patterns of responses. The various eIF4G-Ded1 domain interactions affect *CP-8.1* and *CD-8.1* recruitment quite differently even though these mRNAs differ only by the location of the same SL in an unstructured 5′-UTR. This suggests that there is not a single, uniform mechanism through which Ded1 operates on all mRNAs; instead the diversity of structures in mRNAs requires that Ded1 and the eIF4E·eIF4G·eIF4A·Ded1 complex can operate in multiple modes. The structural diversity inherent in mRNAs presents a challenge for the translational machinery because, once the eIF4F complex attaches to the 5′-cap, structural elements could be oriented in a variety of locations in three-dimensional space relative to its functional domains (Fig. 7C). This problem could explain why eIF4G is so large and flexible and has multiple RNA- and factor-binding domains, which might confer sufficient plasticity to interact with mRNA structures presented in a variety of orientations and distances. Likewise, the multiple Ded1 binding domains on eIF4G might allow Ded1 to assume different positions relative to the diverse mRNA structures it encounters on different mRNAs. The mRNA specificity of effects of the truncation mutants of Ded1 and eIF4G on both K_1/2_ and k_max_ values are consistent with the notion that the eIF4E·eIF4G·eIF4A·Ded1 complex can interact with and modulate the structures of mRNAs in different ways, with the mRNA structure dictating the particular interactions of Ded1 with eIF4G or eIF4A that are most crucial for rapid recruitment.

It is likely that the rate-limiting step(s) for 48S PIC formation will also vary depending on the unique structural features of the mRNA. For some mRNAs, PIC attachment to the 5′-UTR might be rate-limiting because of structures proximal to the 5′-end or because the 5′-end is occluded within the global structure of the mRNA (Fig. 7D (i)). For other mRNAs, PIC attachment might be relatively fast, but scanning to the start codon could be impeded by stable structures that require Ded1 in the context of the eIF4EeIF4GeIF4ADed1 complex to resolve (Fig. 7D (ii)). Our data could be interpreted to suggest that the eIF4G-RNA2 domain is particularly important for Ded1 disruption of cap-distal structures during scanning, as its deletion most strongly reduces k_max_ values for the two mRNAs harboring cap-distal SLs, *SFT2* and *CD-8.1*, with a considerably smaller effect on *SFT2-M* lacking this SL (Fig. 7B). The eIF4G-RNA3/Ded1-CTD interaction could have a role in PIC attachment as removing either of these two domains reduces k_max_ values for mRNAs presumed to have some global structure that might occlude the 5′-end (all mRNAs except *-SL* and *CD-8.1)*, with the largest effects on the two natural mRNAs with short 5′-UTRs that might be sequestered within the global structures created by the bodies of these mRNAs (Fig. 7B, *RPL41A* and *HOR7*). Some mRNAs exhibited recruitment by Ded1 in the absence of eIF4G, including *SFT2, SFT2-M*, and *CD-8.1* (Figs. 3D, 3E, 3G) indicating that Ded1 is also capable of stimulating one or more aspects of 48S PIC assembly outside of the context of the eIF4E·eIF4G·eIF4A·Ded1 complex on certain mRNAs (Fig. 7D (ii)).

These two possible models explaining the differential effects of the eIF4G and Ded1 domain deletions on different mRNAs are not mutually exclusive. In fact, the proposal that the domains have some specificity for mediating PIC attachment versus scanning probably requires that they localize Ded1 to different parts of the mRNA because the former reaction would occur closer to the 5′-end whereas the latter would occur distal to it. The length and flexibility of eIF4G, coupled with the complex network of interactions possible among eIF4G, eIF4E, eIF4A, Ded1 and mRNA, could have evolved to support the plasticity required to deal with the wide variety of mRNA shapes, sizes and structures that must be loaded onto PICs for translation in eukaryotic cells where transcription and translation are uncoupled.

## METHODS

### Preparation of mRNAs and charged initiator tRNA

Plasmids for in vitro run-off mRNA transcription of Ded1-hypodependent and - hyperdependent mRNAs were constructed using Gibson assembly (Gibson et al., 2009). The 5′-UTR and first 60 nucleotides of the coding region of *OST3, SFT2, PMA1, HOR7*, and *FET3* genes were PCR amplified from yeast genomic DNA (BY4741) and cloned into pBluescript II KS+ vector (Stratagene) using NEBuilder HiFi assembly according to the manufacturer’s instructions (New England Biolabs). In all mRNAs constructs, Xmal restriction site was added at the end of the coding region during cloning to linearize the plasmids, and two G nucleotides were added at the beginning of the 5′-UTR to improve transcription efficiency. Plasmids for transcription of *SFT2-M, SFT2-M2*, and *PMA-1M* mRNAs were derived from *SFT2* and *PMA1* plasmids, respectively, by mutating their 5′-UTRs (Genscript Corp.). Plasmids for transcription of *RPL41A*, synthetic mRNAs with 5′-UTR consisting of CAA repeats, and *RPL41A* ORF and 3′-UTR, and initiator tRNA were described previously (Acker et al., 2007; Mitchell et al., 2010; Yourik et al., 2017).

mRNAs and initiator tRNA were transcribed by run-off transcription using T7 RNA polymerase and gel purified as described previously (Acker et al., 2007; Mitchell et al., 2010). mRNAs were capped (m^7^GpppG) using either α-^32^P radiolabeled GTP (Perkin Elmer) or unlabeled GTP and vaccinia virus capping enzyme (Mitchell et al., 2010). Initiator tRNA was methionylated in vitro using methionine and *E. coli* methionyl-tRNA synthetase, and charged Met-tRNA_i_^Met^ was purified of contaminating nucleotides over a desalting column (Walker and Fredrick, 2008; Yourik et al., 2017).

### Purification of translation initiation factors

Eukaryotic initiation factors-eIF1, eIF1A, eIF2, eIF3, eIF4A, eIF4B, eIF4G·4E (WT and mutants), eIF5-were expressed and purified as described previously (Acker et al., 2007; Mitchell et al., 2010; Rajagopal et al., 2012). 40S ribosomal subunits were prepared as described in (Munoz et al., 2017). Recombinant Ded1 proteins (N-terminal Hi_s6_-tag, pET22b vector)-WT Ded1(1-604), Ded1^E307A^, Ded1-ΔCTD (1-535) and Ded1-ΔNTD (117-604) were purified as described previously (Gao et al., 2016; Hilliker et al., 2011) with some modifications. Ded1 proteins were expressed in *E. coli* BL21(DE3) RIL CodonPlus cells (Agilent). Cells were grown at 37°C till OD_600_ of 0.5, cooled to 22°C, and induced with 0.5 mM IPTG overnight. Cells were re-suspended in the lysis buffer (10 mM HEPES-KOH, pH-7.4, 200 mM KCl, 0.1% IGEPAL CA-630, 10 mM imidazole, 10% glycerol, 10 mM 2-mercaptoethanol, DNaseI (1U/ml) and cOmplete protease inhibitor cocktail (Roche)), and lysed using a French Press. Ded1 was purified over a nickel column (5 ml His-Trap column, GE Healthcare) followed by phosphocellulose chromatography (P11, Whartman). Purified protein was dialyzed into dialysis buffer (10 mM HEPES-KOH, pH 7.4, 200 mM KOAc, 50% Glycerol, 2 mM DTT) and stored at −80°C. Ded1-ΔNTD was purified with the same method as the wild-type Ded1 with the following modifications. The N-terminal His6-SUMO tag was removed by incubating nickel-column purified protein with a His6-SUMO protease (McLab) at 4°C overnight, followed by second round of nickel column purification (Gao et al., 2016).

### mRNA recruitment assay

48S PICs were assembled and native gel shift assay was performed as described previously (Mitchell et al., 2010; Yourik et al., 2017) with following modifications. Reactions were assembled in 1X Recon buffer (30 mM HEPES-KOH, pH 7.4, 100 mM KOAc, 3 mM Mg(OAc)2, and 2 mM DTT) containing 300 nM eIF2, 0.5 mM GDPNPMg^2+^, 200 nM Met-tRNAi^Met^, 1 μM eIF1, 1 μM eIF1A, 300 nM eIF5, 300 nM eIF4B, 300 nM eIF3, 30 nM 40S subunits, eIF4A, eIF4EeIFG, and Ded1. The non-hydrolyzable GTP analog GDPNP was used in forming the TC to stabilize the 43S and 48S complexes by preventing conversion to the eIF2·GDP state. The concentrations of eIF4A, eIF4EeIFG, and Ded1 varied for recruitment of different mRNAs. eIF4A: *RPL41A, HOR7, -SL, CP-8.1*, and *CD-8.1* = 7 μM; *SFT2*, *SFT2-M*, *SFT2-M2*, *OST3*, *PMA-1*, *PMA-1M* and *FET3* = 15 μM. eIF4E·eIFG = 75 nM for all mRNAs, except *OST3* (150 nM). Ded1: *RPL41A* and -*SL* = 250 nM, *HOR7* = 500 nM, all other mRNAs = 1 μM. The concentrations of initiation factors were saturating except for the titrant. Reactions were incubated at 26°C for 10 minutes, and were initiated by simultaneous addition of 15 nM ^32^P-m^7^G mRNA and 5 mM ATPMg^2+^. For kinetic analysis, 2 μl aliquot were removed at appropriate times, reactions were stopped by addition of 600-1000 nM non-radiolabeled m^7^G-mRNA (same mRNA as the ^32^P-m^7^G mRNA), and loaded onto a 4% non-denaturing PAGE gel to separate 48S PICs from the free mRNA. Percentage of mRNA recruited to the 48S PIC was calculated using ImageQuant software (GE Healthcare). Data were fitted with a single exponential rate equation to calculate apparent rate of recruitment using KaleidaGraph software (Synergy). Apparent rates were plotted against the concentration of the titrant and fitted with hyperbolic equation to calculate the maximal rates of recruitment and the concentration of the titrant required to achieve the half-maximal rates (K_l/2_). To measure the maximal endpoints of recruitment, the reactions were incubated for 100-200 minutes (as judged by the kinetic experiments). Prism 7 (GraphPad) was used for the statistical analyses and bar-graph data representations. Heatmaps were made in RStudio using gplots and RColorBrewer libraries.

Dissociation experiments were conducted to measure any off-rates (koff) to verify that the addition of unlabeled m^7^G-mRNA to stop the reactions did not result in the dissociation of already-formed 48S PICs. PICs were assembled, and reactions were initiated as described above. A 20-30-fold excess of same unlabeled m^7^G-mRNA was added before the reactions were initiated (no mRNA recruitment was observed) or after 2-20 minutes of incubation (depending on the k_app_). Aliquots were loaded on the gel at indicated times, and percentage mRNA recruited was calculated. The data were fitted with a linear curve.

### ATPase assay

NADH-coupled ATPase assay was performed as described in (Yourik et al., 2017). Ded1 (100-500 nM) was added to reactions containing 1X Recon buffer, 2.5 mM phosphoenolpyruvate, 1 mM NADH, 1/250 dilution of the PK/LDH mix (pyruvate kinase (600 1,0 units/mL) and lactate dehydrogenase (900-1400 units/mL)), 2 μM uncapped *RPL41A* mRNA, and reactions were initiated by addition of 5 mM ATPMg^2+^. Reaction rates (V0) were calculated from linear slope of plot of NADH oxidation over time measured as absorbance A340. k_cat_ was calculated by dividing V0 by Ded1 concentration. To estimate the K_m_ of ATP, ATP was titrated at 0-10 mM concentrations. Rates were plotted against the concentration of the ATP and fitted with Michaelis-Menten equation to calculate the K_m_ of ATP.

### Fluorescence anisotropy assay

Fluorescent anisotropy assay was performed as described previously (Walker et al., 2013). Briefly, ssRNA labeled with fluorescein at the 3′-end was incubated with Ded1 concentration (0-750 nM) in the absence of any nucleotide or in the presence of 5 mM ADPMg^2+^ and ADPNPMg^2+^ in 1X Recon buffer, and fluorescent anisotropies were measured with excitation and emission wavelengths of 495 nM and 520 nM, respectively. The data were fitted with a hyperbolic equation.

## ACKNOWLEDGEMENTS

We thank Roy Parker and Eckhard Jankowsky for gifts of Ded1 expression plasmids, and Thomas Dever and Nicholas Guydosh for thoughtful comments and suggestions.

## List of supplementary figures

**Figure S1 related to Figure 1:**
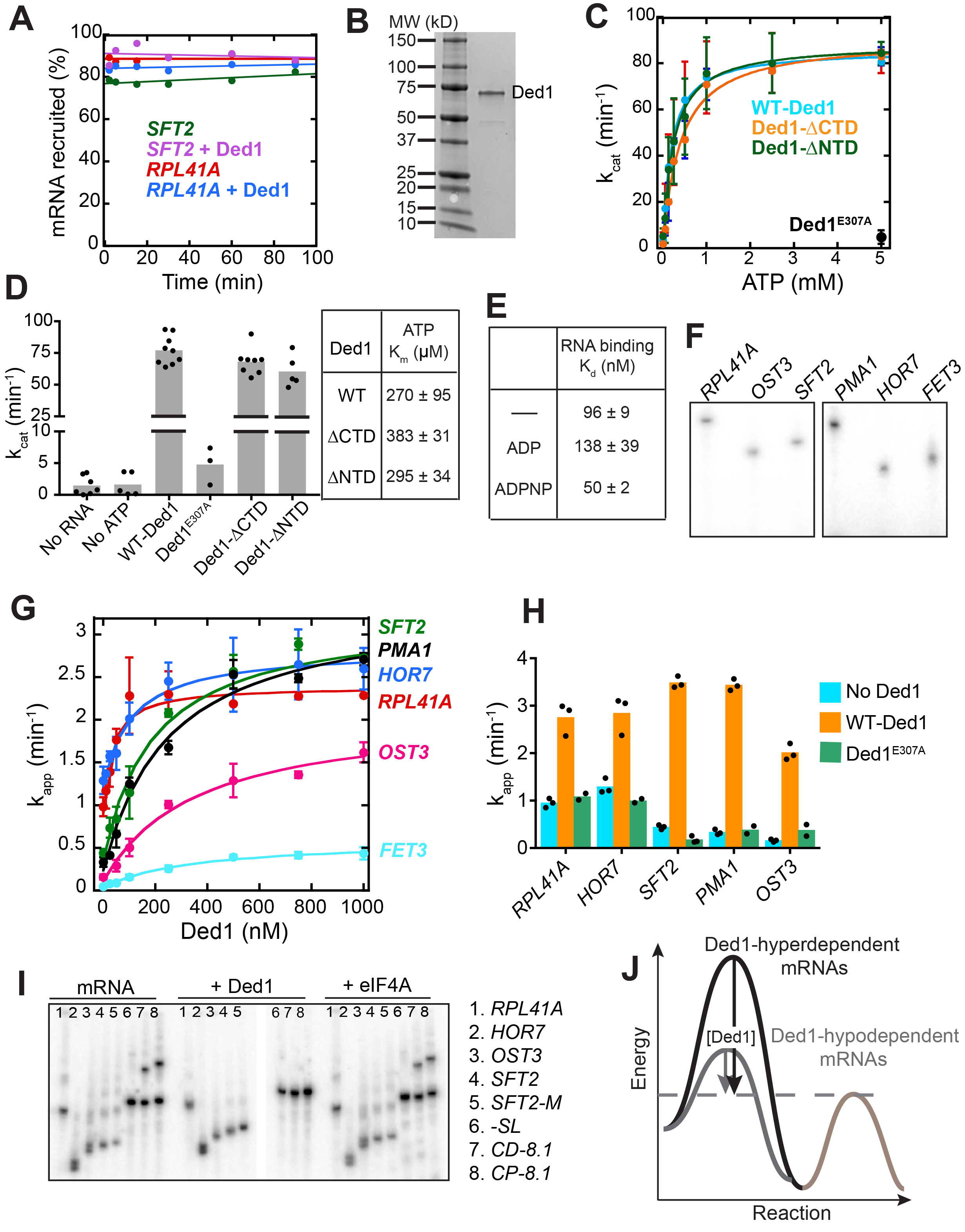
Catalytically active Ded1 stimulates 48S PIC assembly on native mRNAs. (A) Dissociation of 48S PICs assembled on *RPL41A* and *SFT2* mRNAs after addition of excess cold (non-radiolabeled) capped mRNA as a “pulse-quench” was monitored over time. 48S PICs were assembled for 15 min with or without Ded1 and chased with an ~20-fold molar excess of the corresponding cold-capped mRNAs. The percentage of labeled mRNA remaining bound to 48S PICs was quantified after the indicated times of “pulse-quench”. No detectable dissociation was observed over the indicated time-course for these two mRNAs, or for any other mRNAs depicted in Figure 1A (data not shown), indicating that 48S PICs assembled on *RPL41A* and *SFT2* mRNAs are highly stable and that cold-capped mRNAs only sequester unbound PICs. **(B)** Purified Hi_s6_-Ded1 and size markers (lane 1) were resolved by SDS-PAGE on a 4-15% gel and stained with SimplyBlue Safestain. **(C)** Observed rates of RNA-dependent ATP hydrolysis at different ATP concentrations in the presence of saturating (2 μM) uncapped *RPL41A* mRNA for WT and mutant Ded1 proteins, the latter harboring the alanine substitution of Glu-307 (E307A) in the DEAD motif (Ded1^E307A^); or truncations of the NTD or CTD domain, as indicated in Figure S4A. Data from two independent experiments are shown, plotting mean values ± 1 SD. Data was fitted with the Michaelis-Menten equation. **(D)** k_cat_ values (at saturating ATP concentration of 5 mM) and 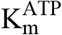 were determined from the experiments described in (C). WT Ded1, Ded1-ΔCTD, and Ded1-ΔNTD have similar k_cat_ and 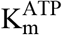 values, whereas reaction kinetics with Ded1^E307A^ are comparable to “No mRNA” and “No ATP” controls conducted with WT Ded1. The ATPase activity of WT Ded1 is similar to previously published values, ranging from k_cat_ = 30-680 min^−1^ depending on the RNA substrate used, and K_m_ values for ATP of 280340 μ.M (Iost et al., 1999; Senissar et al., 2014). Note the discontinuity in the Y-axis. Bars represent the mean with dots as individual experimental values (n ≥ 3). 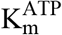 is expressed as mean values ± 1 SD. **(E)** Binding affinity of WT Ded1 for single-stranded RNA was determined in the presence or absence of the indicated nucleotides (5 mM) by measuring changes in fluorescence anisotropy of a single-stranded RNA labeled with 3′-6-Fluorescein over Ded1 concentrations ranging from 0 to 750 nM. The determined K_d_ values are similar to previously reported measurements: K_d_ = 180-230 nM (no nucleotide); 210-290 nM (ADP); and 35-40 nM (ADPNP) (Banroques et al., 2008; Iost et al., 1999). Addition of buffer in the absence of Ded1 produced no change in fluorescence anisotropy of the RNA. Mean values ± 1 SD are reported (n=2). **(F)** [α^32^P]-capped mRNAs used in mRNA recruitment assays in Fig. 1A resolved by electrophoresis on a 6% denaturing urea-TBE gel. **(G)** The apparent rates (k_app_) of 48S PIC assembly measured as a function of Ded1 concentration for the indicated mRNAs. Data were fitted to a hyperbolic curve with an offset to account for the rates observed in the absence of Ded1, allowing calculation of k_max_ and 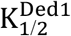 values reported in Figs. 1B-C, respectively. Data points are mean values with error bars (1 SD) at different Ded1 concentrations (n = 3). **(H)** Ded1 stimulation of mRNA recruitment is dependent on the ATPase activity of Ded1. Mean observed rates of 48S assembly (n=3) with no Ded1 (blue bars), saturating Ded1 (orange bars, determined as in Fig. 1B), and ATPase deficient mutant Ded1^E307A^ (green bars). **(I)** [α^32^P]-capped mRNAs resolved by electrophoresis on a 4% non-denaturing gel. mRNAs exhibit isoforms of different mobility (mRNA only lanes). Ded1 resolved most mRNA conformations to a single conformation (+Ded1 lanes) whereas eIF4A had a negligible effect (+eIF4A lanes). The indicated mRNAs (15 nM) were incubated for 10 min with 5 mM ATP and either 100 nM Ded1 or 2 μ.M eIF4A. **(J)** Ded1-hyperdendent mRNAs have a higher energy barrier (dark grey) as compared to Ded1-hypodependent mRNAs (light grey) and require higher Ded1 concentrations to lower the activation energy for the Ded1-accelerated rate-limiting step until the energy barrier for a non-Ded1 dependent step (blue) becomes rate-limiting. For *OST3* and *FET3* mRNAs, Ded1 can only partially lower the energy barrier (red dotted line). See SuppFig1_SourceData1

**Figure S2 related to Figures 1 and 2:**
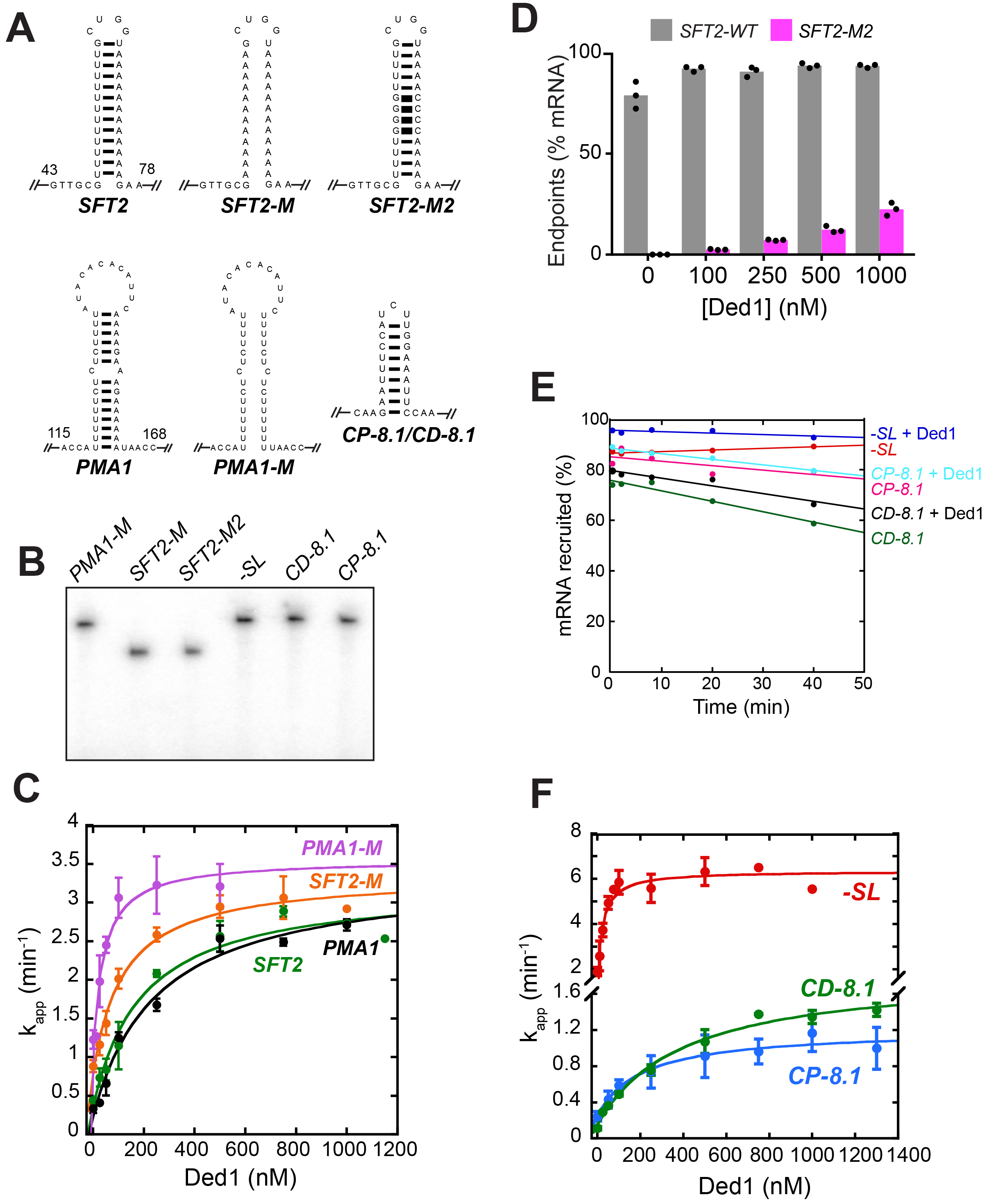
Ded1 enhances the recruitment of mRNAs with SLs in the 5′-UTRs. (A) Sequence and structures of SLs in the 5′-UTRs of mRNAs examined in this study (Figures 1A and 2A) of predicted ΔG°s: *SFT2* = −9.4 kcal/mol, *SFT2-M2* = −19.7 kcal/mol, *PMA1* = −8.2 kcal/mol, *CP-8.1/CD-8.1* = −8.1 kcal/mol. The distance from the 5′-end is indicated for *SFT2* and *PMA1.* (B) [α^32^P]-capped mRNAs used in 48S PIC assembly assays, depicted in Fig. 2A, resolved by electrophoresis on a 6% denaturing urea-TBE gel. **(C)** The k_app_ for mRNA recruitment measured as a function of Ded1 concentrations for *PMA1* (black), *PMA1-M* (purple), *SFT2* (green), and *SFT2-M* (orange) mRNAs. Data were fitted with a hyperbolic curve with an offset to account for the rate observed with no Ded1, allowing calculation of k_max_ and 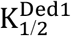 values reported in Figs. 2C and 2D, respectively. Plotted points are mean values with error bars indicating 1 SD (n = 3). **(D)** Endpoints of 48S PIC assembly at different Ded1 concentrations for *SFT2* and *SFT2-M2* mRNAs. The endpoints for *SFT2-M2* represent lower limits because Ded1 concentrations above 1 μM were unattainable. **(E)** *-SL, CP-8.1*, and *CD-8.1* mRNAs assembled in 48S PICs (+/− Ded1) remain stably bound for 40 min after addition of ~20-fold cold-capped (non-radiolabeled) mRNA to stop the reactions, conducted as described in Fig. S1A. Modest dissociation was observed for *CP-8.1* and *CD-8.1* mRNAs; but none for *-SL* mRNA. **(F)** The k_app_ for mRNA recruitment measured as a function of Ded1 concentration for *-SL* (red), *CP-8.1* (blue), and *CD-8.1* (green) mRNAs. The data were fitted with a hyperbolic curve with an offset to account for the rate observed with no Ded1, allowing calculation of k_max_ and 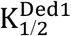 values reported in Figs. 2C and 2D, respectively. Plotted points are the mean values with error bars of 1 SD (n = 3). Note the y-axis is discontinuous. See SuppFig2_SourceData1

**Figure S3 related to Figure 3:**
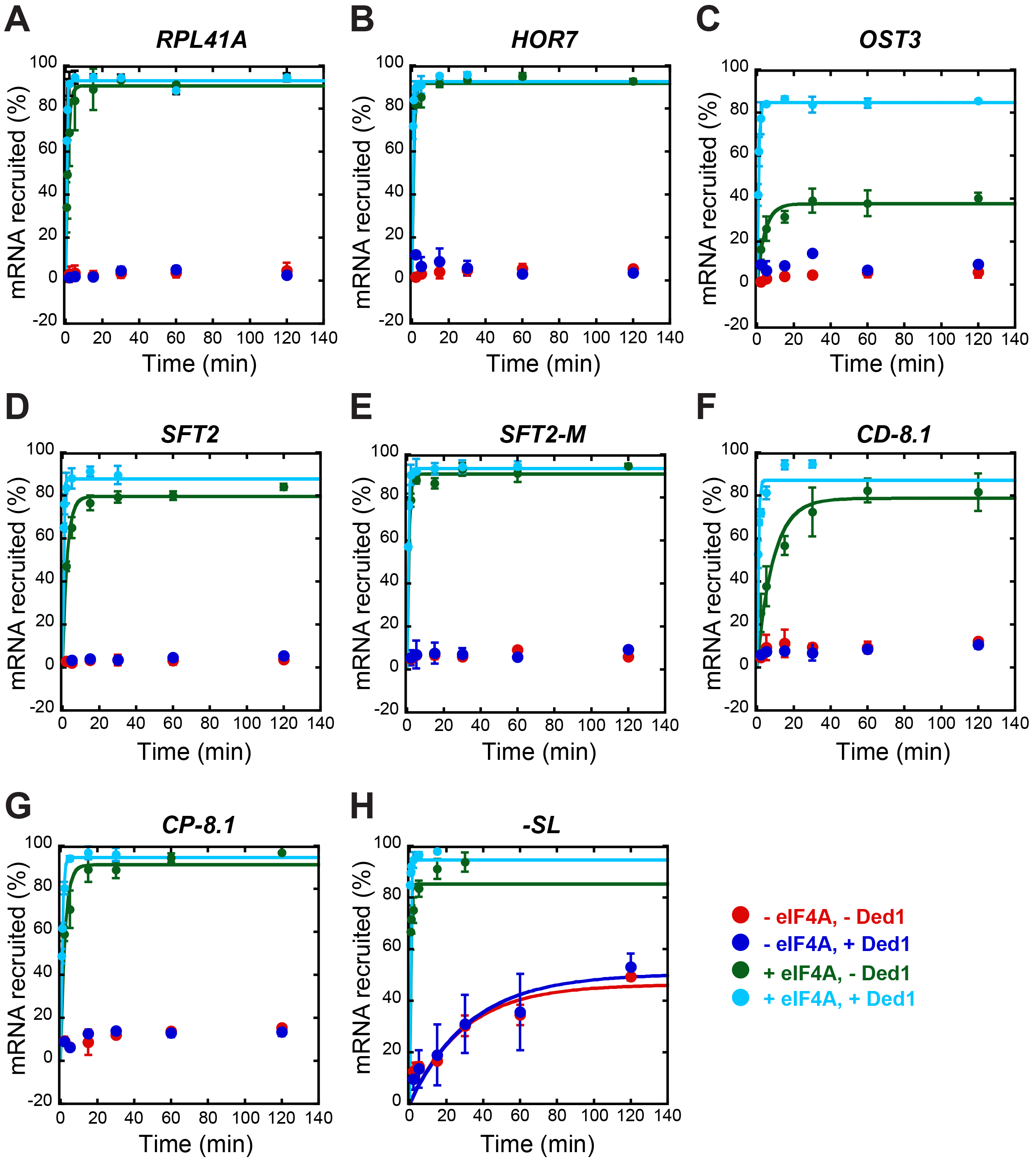
eIF4A is required for appreciable recruitment of all mRNAs tested (except *-SL*) in the presence or absence of saturating Ded1. Recruitment of mRNAs without eIF4A in the absence (red) or presence of Ded1 (blue), and with eIF4A in the absence (green) or presence of Ded1 (cyan). The data were fitted with a hyperbolic curve for the reactions including eIF4A in the absence (green) and presence of Ded1 (cyan); and for all four conditions for *-SL* mRNA (H). Each point represents the mean value (n ≥ 2) and error bars are 1 SD from the mean. See SuppFig3_SourceData1

**Figure S4 related to Figures 4, 5 and 6:**
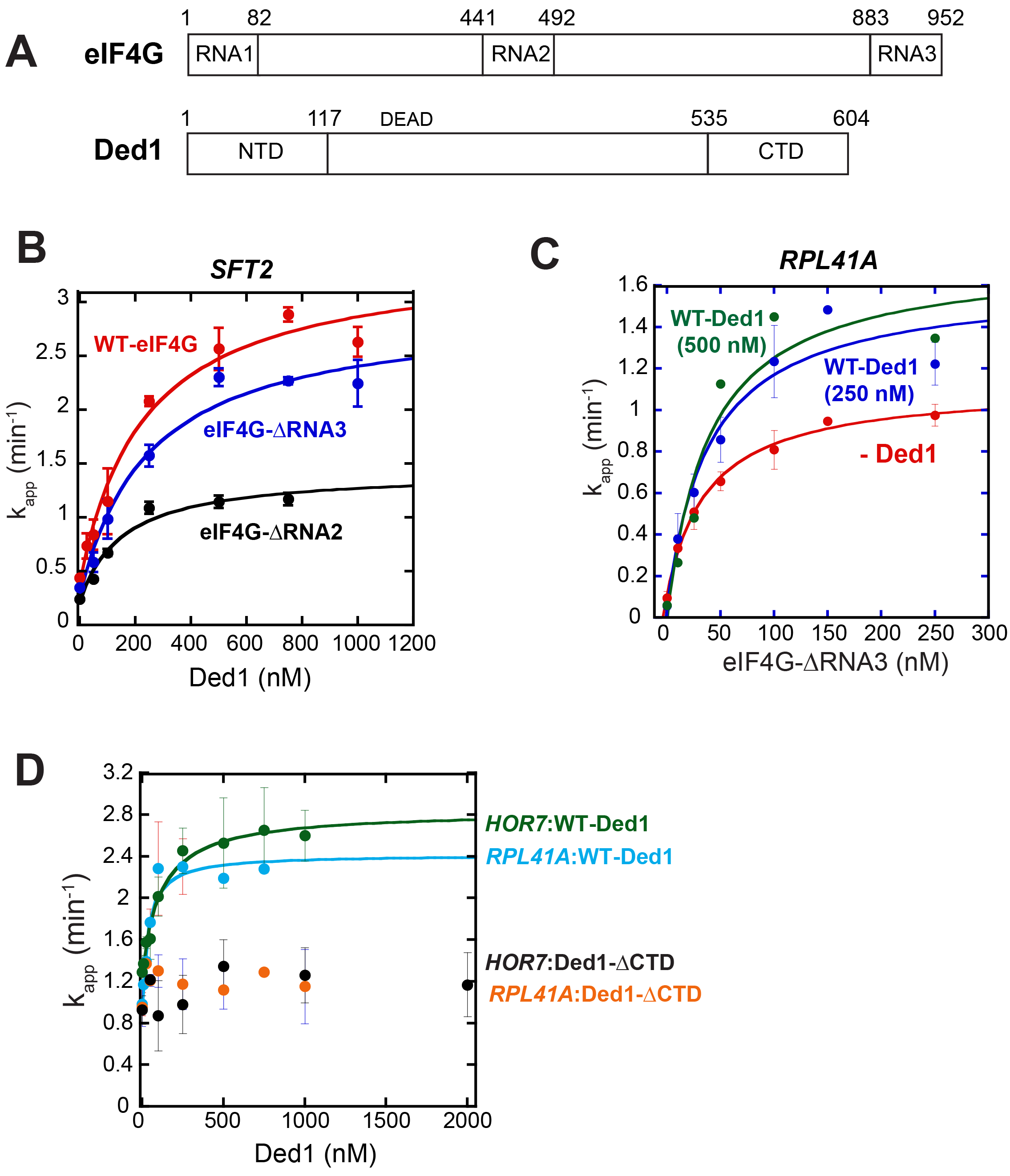
Selected effects of the eIF4G-ΔRNA3 and Ded1-ΔCTD truncations on recruitment of native mRNAs. **(A)** Schematics of eIF4G showing RNA binding domains-RNA1, RNA2, and RNA3 and Ded1 showing N-terminal and C-terminal domains, indicating the amino acid coordinates at domain junctions. **(B)** The apparent rate of *SFT2* mRNA recruitment measured as a function of Ded1 concentration with WT eIF4G (75 nM), eIF4G-ΔRNA3 (150 nM) and eIF4G-ΔRNA2 (400 nM). Data fitted with a hyperbolic curve. Plotted points are mean values with error bars of 1 SD (n = 3). The determined 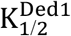 values are 236 ± 18 nM, 251 ± 34 nM, 148 ± 17 nM, with WT eIF4G, eIF4G-ΔRNA3, and eIF4G-ΔRNA2, respectively; indicating that the Ded1 concentrations used in Figs. 4 and 5 were saturating for *SFT2* mRNA. Similar experiments were also conducted for the other mRNAs examined in Figs. 4 and 5 using a range of Ded1 concentrations up to 1μM, which corresponds to 2X-4X of the 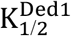 obtained with WT eIF4G in Figs. 1C and 2D. The results (data not shown) ensured that Ded1 was present at saturating levels in the reactions for these mRNAs containing the two eIF4G mutants. **(C)** Recruitment of *RPL41A* as a function of eIF4G-ΔRNA3 concentration with no Ded1 (red), 250 nM Ded1 (blue) and 500 nM Ded1 (green). The results indicate that doubling the Ded1 concentration from that employed in Fig. 4C did not increase the rate or change the 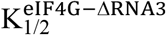 of 48S PIC assembly. **(D)** The apparent rates of *RPL41A* and *HOR7* recruitment observed at different concentrations of WT Ded1 and Ded1-ΔCTD. Data obtained with WT Ded1 were fitted with a hyperbolic curve. The results indicate a negligible change in rate using Ded1-ΔCTD. See SuppFig4_SourceData1

**Figure S5 related to Figures 4, 5 and 6:**
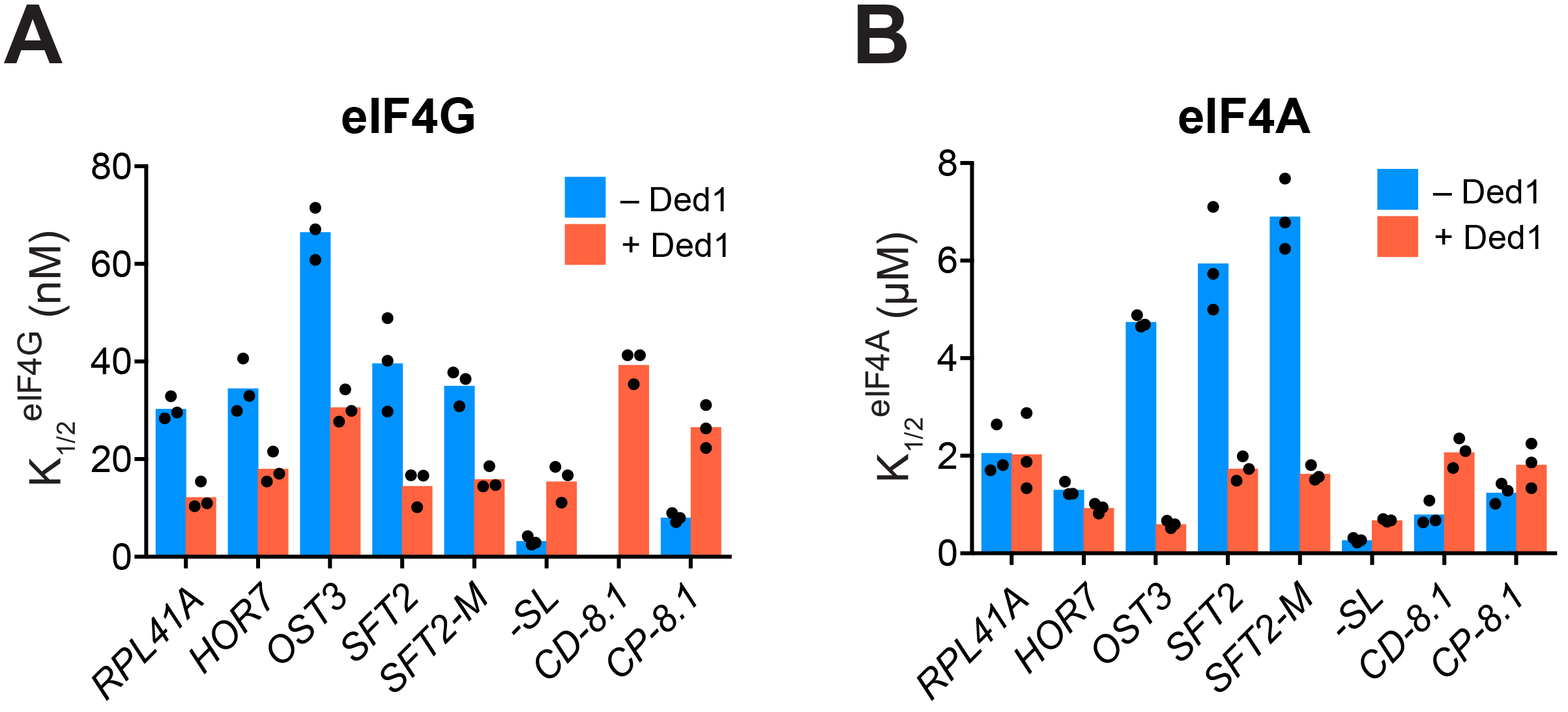
**(A, B)** K_1/2_ values of eIF4G·4E (A) and eIF4A (B) in the absence of Ded1 (dark blue) and presence of saturating Ded1 (red). Bars indicate mean values calculated from 3 independent experiments represented by the data points. We probed functional interactions between Ded1 and eIF4G by measuring the apparent recruitment rates at varying concentrations of eIF4G in the presence and absence of Ded1. In the absence of Ded1, the 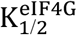 for mRNAs *RPL41A, HOR7, OST3, SFT2*, and *SFT2-M*ranged from 30 to 75 nM (panel A, cols. 1-5, blue); whereas the presence of saturating Ded1 decreased the 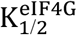 by factors of ~2-to 2.5 for each of these five mRNAs (red vs. blue). The reductions in 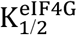 conferred by Ded1 are consistent with the idea that Ded1 interacts productively with eIF4G for all five of these mRNAs. Similarly, the presence of saturating Ded1 lowered the concentration of eIF4A required to achieve the half-maximal rate of recruitment 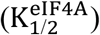 by 3-7-fold for *OST3, SFT2*, and *SFT2-M* (panel B, cols. 3-5, red vs. blue)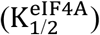, consistent with a productive interaction between Ded1 and eIF4A during 48S formation on these mRNAs. This interaction could be direct, through eIF4G, or due to Ded1 altering the conformation of the mRNA or another component of the system, making it easier for eIF4A to bind or act on it. The native mRNAs *RPL41A* and *HOR7*, has the same 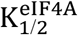 with and without Ded1 (panel B, cols. 1-2, red vs. blue). As the 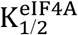 for *RPL41A* and *HOR7* in the absence of Ded1 were lower than those for *OST3, SFT2*, and *SFT2-M* (panel B, blue), and were actually at the level reached for *SFT2* and *SFT2-M* in the presence of Ded1 (panel B, red), perhaps *RPL41A* and *HOR7* are naturally more receptive to the effects of eIF4A without assistance from Ded1, possibly because of their short 5′-UTR. The synthetic mRNAs produced distinctly different functional interactions between Ded1 and eIF4G or eIF4A. In the absence of Ded1, *-SL* and *CP-8.1* had lower 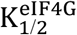 values of 3 to 8 nM compared to 30-75 nM for the five natural mRNAs (panel A, blue bars). (The 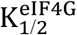 for *CD-8.1* without Ded1 could not be determined accurately because of its endpoint defects at lower eIF4G concentrations.) Contrary to the natural mRNAs, addition of Ded1 increased the 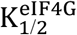 of *-SL* and *CP-8.1* mRNAs by 3-5-fold, reaching values similar to those observed for the natural mRNAs in the presence of Ded1 (panel A, red bars). The synthetic mRNAs also exhibited 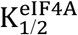 values in the absence of Ded1 that were generally lower than the corresponding values for the native mRNAs (panel B, blue bars). Moreover, addition of Ded1 increased the 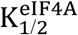 values of the two synthetic mRNAs, in contrast to the decreased 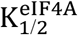 values evoked by Ded1 for three of the five native mRNAs (panel A, red). These results indicate that the maximum stimulation of recruitment of the synthetic mRNAs by eIF4G and eIF4A in the absence of Ded1 can be achieved at relatively low concentrations of both factors, but that higher concentrations of eIF4G and eIF4A are required to support the additional stimulation of recruitment conferred by Ded1, which are comparable in magnitude to the concentrations of eIF4G and eIF4A required for maximal Ded1-stimulation of the native mRNAs. In summary, the requirements for eIF4G and eIF4A, as well as the effect of Ded1 on these requirements, differ among different native mRNAs and vary dramatically between native and synthetic mRNAs. These results suggest the existence of different functional modalities of the eIF4F complex (in the absence of Ded1) and of the eIF4E-eIF4G-eIF4A-Ded1 complex (with Ded1) that may be required to overcome the range of RNA structures in different mRNAs that vary in stability, distance from the 5′-cap, or orientation in three-dimensional space to accelerate mRNA attachment and subsequent scanning to the AUG codon. (See Discussion for additional comments.) See SuppFig5_SourceData1

## REFERENCES

Abaeva, I.S., Marintchev, A., Pisareva, V.P., Hellen, C.U., and Pestova, T.V. (2011). Bypassing of stems versus linear base-by-base inspection of mammalian mRNAs during ribosomal scanning. EMBO J 30, 115–129.

Acker, M.G., Kolitz, S.E., Mitchell, S.F., Nanda, J.S., and Lorsch, J.R. (2007). Reconstitution of yeast translation initiation. Methods Enzymol 430, 111–145.

Banroques, J., Cordin, O., Doere, M., Linder, P., and Tanner, N.K. (2008). A conserved phenylalanine of motif IV in superfamily 2 helicases is required for cooperative, ATP-dependent binding of RNA substrates in DEAD-box proteins. Mol Cell Biol 28, 3359–3371.

Berset, C., Zurbriggen, A., Djafarzadeh, S., Altmann, M., and Trachsel, H. (2003). RNA-binding activity of translation initiation factor eIF4G1 from Saccharomyces cerevisiae. RNA 9, 871–880.

Berthelot, K., Muldoon, M., Rajkowitsch, L., Hughes, J., and McCarthy, J.E. (2004). Dynamics and processivity of 40S ribosome scanning on mRNA in yeast. Mol Microbiol 51, 987–l00l.

Bowers, H.A., Maroney, P.A., Fairman, M.E., Kastner, B., Luhrmann, R., Nilsen, T.W., and Jankowsky, E. (2006). Discriminatory RNP remodeling by the DEAD-box protein Ded1. RNA 12, 903–912.

Chiu, W.L., Wagner, S., Herrmannova, A., Burela, L., Zhang, F., Saini, A.K., Valasek, L., and Hinnebusch, A.G. (2010). The C-terminal region of eukaryotic translation initiation factor 3 a (eIF3a) promotes mRNA recruitment, scanning, and, together with eIF3j and the eIF3b RNA recognition motif, selection of AUG start codons. Mol Cell Biol 30, 4415–4434.

Chuang, R.Y., Weaver, P.L., Liu, Z., and Chang, T.H. (1997). Requirement of the DEAD-Box protein Ded1p for messenger RNA translation. Science 275, 1468–1471.

de la Cruz, J., Iost, I., Kressler, D., and Linder, P. (1997). The p20 and Ded1 proteins have antagonistic roles in eIF4E-dependent translation in Saccharomyces cerevisiae. Proc Natl Acad Sci U S A 94, 5201–5206.

Dever, T.E., Kinzy, T.G., and Pavitt, G.D. (2016). Mechanism and Regulation of Protein Synthesis in Saccharomyces cerevisiae. Genetics 203, 65–107.

Gao, Z., Putnam, A.A., Bowers, H.A., Guenther, U.P., Ye, X., Kindsfather, A., Hilliker, A.K., and Jankowsky, E. (2016). Coupling between the DEAD-box RNA helicases Ded1p and eIF4A. Elife 5.

Gibson, D.G., Young, L., Chuang, R.Y., Venter, J.C., Hutchison, C.A. 3rd,, and Smith, H.O. (2009). Enzymatic assembly of DNA molecules up to several hundred kilobases. Nat Methods 6, 343–345.

Hilliker, A., Gao, Z., Jankowsky, E., and Parker, R. (2011). The DEAD-box protein Ded1 modulates translation by the formation and resolution of an eIF4F-mRNA complex. Mol Cell 43, 962–972.

Hinnebusch, A.G. (2014). The scanning mechanism of eukaryotic translation initiation. Annu Rev Biochem 83, 779–812.

Iost, I., Dreyfus, M., and Linder, P. (1999). Ded1p, a DEAD-box protein required for translation initiation in Saccharomyces cerevisiae, is an RNA helicase. J Biol Chem 274, 17677–17683.

Kertesz, M., Wan, Y., Mazor, E., Rinn, J.L., Nutter, R.C., Chang, H.Y., and Segal, E. (2010). Genome-wide measurement of RNA secondary structure in yeast. Nature 467, 103–107.

Kumar, P., Hellen, C.U., and Pestova, T.V. (2016). Toward the mechanism of eIF4F-mediated ribosomal attachment to mammalian capped mRNAs. Genes Dev 30, 1573–1588.

Linder, P., and Slonimski, P.P. (1989). An essential yeast protein, encoded by duplicated genes TIFl and TIF2 and homologous to the mammalian translation initiation factor eIF-4A, can suppress a mitochondrial missense mutation. Proc Natl Acad Sci U S A 86, 2286–2290.

Mitchell, S.F., Walker, S.E., Algire, M.A., Park, E.H., Hinnebusch, A.G., and Lorsch, J.R. (2010). The 5′-7-methylguanosine cap on eukaryotic mRNAs serves both to stimulate canonical translation initiation and to block an alternative pathway. Mol Cell 39, 950–962.

Munoz, A.M., Yourik, P., Rajagopal, V., Nanda, J.S., Lorsch, J.R., and Walker, S.E. (2017). Active yeast ribosome preparation using monolithic anion exchange chromatography. RNA Biol 14, 188–196.

Park, E.H., Walker, S.E., Lee, J.M., Rothenburg, S., Lorsch, J.R., and Hinnebusch, A.G. (2011). Multiple elements in the eIF4G1 N-terminus promote assembly of eIF4G1*PABP mRNPs in vivo. EMBO J 30, 302–316.

Pestova, T.V., and Kolupaeva, V.G. (2002). The roles of individual eukaryotic translation initiation factors in ribosomal scanning and initiation codon selection. Genes Dev 16, 2906–2922.

Pisareva, V.P., Pisarev, A.V., Komar, A.A., Hellen, C.U., and Pestova, T.V. (2008). Translation initiation on mammalian mRNAs with structured 5′UTRs requires DExH-box protein DHX29. Cell 135, l237–l250.

Putnam, A.A., Gao, Z., Liu, F., Jia, H., Yang, Q., and Jankowsky, E. (2015). Division of Labor in an Oligomer of the DEAD-Box RNA Helicase Ded1p. Mol Cell 59, 541–552.

Rajagopal, V., Park, E.H., Hinnebusch, A.G., and Lorsch, J.R. (2012). Specific domains in yeast translation initiation factor eIF4G strongly bias RNA unwinding activity of the eIF4F complex toward duplexes with 5′-overhangs. J Biol Chem 287, 20301–20312.

Rogers, G.W. Jr.,, Richter, N.J., and Merrick, W.C. (1999). Biochemical and kinetic characterization of the RNA helicase activity of eukaryotic initiation factor 4A. J Biol Chem 274, 12236–12244.

Rouskin, S., Zubradt, M., Washietl, S., Kellis, M., and Weissman, J.S. (2014). Genome-wide probing of RNA structure reveals active unfolding of mRNA structures in vivo. Nature 505, 701–705.

Sen, N.D., Zhou, F., Ingolia, N.T., and Hinnebusch, A.G. (2015). Genome-wide analysis of translational efficiency reveals distinct but overlapping functions of yeast DEAD-box RNA helicases Ded1 and eIF4A. Genome Res 25, 1196–1205.

Senissar, M., Le Saux, A., Belgareh-Touze, N., Adam, C., Banroques, J., and Tanner, N.K. (2014). The DEAD-box helicase Ded1 from yeast is an mRNP cap-associated protein that shuttles between the cytoplasm and nucleus. Nucleic Acids Res 42, 10005–10022.

Sokabe, M., and Fraser, C.S. (2017). A helicase-independent activity of eIF4A in promoting mRNA recruitment to the human ribosome. Proc Natl Acad Sci U S A 114, 6304–6309.

Walker, S.E., and Fredrick, K. (2008). Preparation and evaluation of acylated tRNAs. Methods 44, 81–86.

Walker, S.E., Zhou, F., Mitchell, S.F., Larson, V.S., Valasek, L., Hinnebusch, A.G., and Lorsch, J.R. (2013). Yeast eIF4B binds to the head of the 40S ribosomal subunit and promotes mRNA recruitment through its N-terminal and internal repeat domains. RNA 19, 191–207.

Yang, Q., Fairman, M.E., and Jankowsky, E. (2007). DEAD-box-protein-assisted RNA structure conversion towards and against thermodynamic equilibrium values. J Mol Biol 368, 1087–1100.

Yang, Q., and Jankowsky, E. (2006). The DEAD-box protein Ded1 unwinds RNA duplexes by a mode distinct from translocating helicases. Nat Struct Mol Biol 13, 981–986.

Yourik, P., Aitken, C.E., Zhou, F., Gupta, N., Hinnebusch, A.G., and Lorsch, J.R. (2017). Yeast eIF4A enhances recruitment of mRNAs regardless of their structural complexity. Elife 6.

